# MaskGraphene: an advanced framework for interpretable joint representation for multi-slice, multi-condition spatial transcriptomics

**DOI:** 10.1101/2024.02.21.581387

**Authors:** Yunfei Hu, Zhenhan Lin, Manfei Xie, Weiman Yuan, Yikang Li, Mingxing Rao, Yichen Henry Liu, Wenjun Shen, Lu Zhang, Xin Maizie Zhou

**Affiliations:** Department of Computer Science, Vanderbilt University, Nashville, USA; Department of Biomedical Engineering, Vanderbilt University, Nashville, USA; Department of Bioinformatics, Shantou University Medical College, Shantou, China; Department of Computer Science, Hongkong Baptist University, Kowloon Tong, Hong Kong

**Keywords:** Spatial Transcriptomics, Batch correction, Integration, Interpretable embeddings, Self- supervised learning, Contrastive learning

## Abstract

Recent advancements in spatial transcriptomics (ST) have underscored the importance of integrating data from multiple ST slices for joint analysis. A major challenge remains generating interpretable joint embeddings that preserve geometric information for downstream analyses. Here we introduce MaskGraphene, a graph neural network that combines self-supervised and self-contrastive training to integrate gene expression and spatial location into joint embeddings. By employing clusterwise alignment and a graph attention autoencoder with masked self-supervised and triplet loss optimizations, MaskGraphene effectively preserves geometric structures while achieving batch correction. In benchmarks against seven state-of-the-art methods, MaskGraphene consistently demonstrated superior alignment accuracy and geometric fidelity across diverse ST datasets. Its interpretable embeddings significantly enhanced downstream applications, including domain identification, spatial trajectory reconstruction, biomarker discovery, and the creation of topographical maps of brain slices. Notably, MaskGraphene successfully recovered layer-wise brain structures with near-perfect accuracy. MaskGraphene provides a powerful and versatile framework for advancing ST data integration and analysis, unlocking valuable biological insights.

## Introduction

The intricate orchestration of biological processes within organisms hinges upon the diversity and specialization of individual cell types, each tailored to fulfill specific functions. To unravel the complexities of disease pathology and tissue functioning, understanding the connections and interactions of neighboring and distant cells is crucial since the behavior of these cells is profoundly influenced by their microenvironment.^1^ While single-cell RNA sequencing (scRNA-seq) has revolutionized our ability to profile gene expression at single-cell resolution, its lack of spatial context limits our understanding of cellular niches within their native microenvironments. This limitation impedes our understanding of key processes such as cell-cell interactions, tissue organization, and spatially regulated functional dynamics, all of which are vital for deciphering biological systems.^2, 3^

Advancements in Spatial Transcriptomics (ST) have bridged the gap between transcriptomic profiling and spatial context by enabling the simultaneous measurement of mRNA expression and spatial coordinates within tissue sections.^4^ These techniques have significantly advanced our ability to explore the complex transcriptional landscapes of heterogeneous tissues.^5^ These methods can be broadly categorized into imaging-based techniques, such as smFISH, STARmap, MERFISH, seqFISH, and seqFISH+, which offer high spatial resolution, and sequencing-based techniques, including Slide-seq, Slide-seqV2, 10x Visium, spatial transcriptomics (ST), HDST, and Stereo-seq, which provide scalable data with varying resolution.^6–10^ Despite these advancements, the integrated analysis of ST datasets generated under different experimental conditions or technologies remains a significant challenge.^11^

Conventional single-slice ST data analyses primarily focus on uncovering spatial domain distributions within individual tissue sections.^12, 3^ Recently, there has been growing recognition of the value of integrative and comparative analysis of ST datasets from diverse sources, including different samples, biological conditions, technological platforms, and developmental stages.^14^ Integrative analysis provides a more comprehensive understanding of spatial tissue structures, enabling deeper insights into complex spatial organization and leveraging additional information for more robust analyses. Nevertheless, ST datasets are susceptible to batch effects, which can obscure true biological signals and complicate data interpretation. Therefore, developing a well-designed multi-slice ST data integration framework that jointly models and harmonizes signals across samples while mitigating batch effects is an urgent and critical need.

Existing integration strategies often focus on aggregating gene expression data across slices or applying batch correction techniques originally developed for scRNA-seq, like Harmony^15^ and Seurat.^16^ These approaches often neglect the vital spatial coordinates of ST data. To address these limitations, advanced tools like BASS,^17^ DeepST,^14^ PRECAST,^18^ GraphST,^19^ SPIRAL,^20^ STAligner,^21^ and SpaDo^22^ have been developed. These methods employ a wide variety of strategies. BASS applies a hierarchical Bayesian model framework for multi-slice clustering, producing clustering labels as outputs. DeepST utilizes a variational graph autoencoder combined with data augmentation techniques to enable joint analysis of ST slices. PRECAST leverages a Gaussian mixture model and hidden Markov random fields to derive latent joint embeddings. GraphST, recognized for its strong performance in single-slice clustering tasks, extends its capabilities to multi-slice analysis by constructing positive and negative spot pairs for contrastive training using representations of both normal and corrupted graphs. SPIRAL employs a graph autoencoder with an optimal transport-based discriminator and a classifier to mitigate batch effects, align spatial coordinates, and refine gene expression. STAligner integrates a graph attention autoencoder with a triplet loss strategy to enhance spatial alignment and embedding quality. SpaDo calculates a unified Spatially Adjacent Cell Type Embedding (SPACE) for multiple slices. This is achieved by concatenating individual slice embeddings and employing Jensen–Shannon divergence and Manhattan distance for hierarchical clustering, ensuring cross-slice comparability through consistent cell type annotations.

With the exception of BASS, all of these methods focus on learning joint embeddings for integration analysis. While they have shown varying degrees of effectiveness in multi-slice analyses,^11^ they share a common limitation inherent to many graph-based deep learning models: the inability to generate interpretable joint embeddings that accurately capture geometric information. This shortfall hinders precise node-wise (or spot-wise) alignment across slices, ultimately limiting their effectiveness in ST data integration and other downstream analyses.

To address the challenges of integrating ST data with interpretable, batch-corrected joint embeddings, we introduce MaskGraphene, a graph neural network that leverages self-supervised and self-contrastive training strategies. MaskGraphene integrates gene expression and spatial context data to generate joint embeddings suitable for integration and various downstream analyses. To ensure interpretable joint embeddings that preserve original geometric information, the model incorporates two types of interslice connections: “hard-links”, which establish direct connections across slices using k-NN networks based on spot-wise alignment, and “soft-links”, which create indirect connections using triplets constructed across slices. Furthermore, its masked self-supervised loss enforces reconstruction in the embedding space while regularizing the node feature reconstruction, enhancing both interpretability and robustness.

We benchmarked MaskGraphene against seven integration methods across a diverse range of spatial transcriptomics datasets, evaluating performance for various integration scenarios and quantitative metrics. MaskGraphene consistently achieved the highest alignment accuracy and matching performance among all tools. Notably, its joint embeddings, visualized in two-dimensional UMAP, are the first to effectively capture the overall slice shape, partially overlapping patterns, spatial structures, and geometric relationships. Taking advantage of its interpretable joint embeddings, MaskGraphene excelled in multiple integration scenarios and downstream analyses, including enhanced domain identification, reconstruction of spatial trajectories across slices, improved biomarker identification, generation of topographical maps of brain slices, identification of neuronal differentiation and activity gradients, simultaneous characterization of unique and shared tissue structures across horizontally consecutive slices, and alignment of tissue structures across developmental stages.

## Results

### Overview of MaskGraphene

In the workflow (Fig.1), MaskGraphene processes a spot-gene expression matrix and spatial coordinates from two or more tissue slices, tailored to various integration scenarios. To enhance interslice connectivity, cluster-wise alignment is first performed for each pair of consecutive slices to align spots across slices, establishing direct “hard-links”, which are then used to construct a k-NN graph for the graph model. MaskGraphene adopts specific strategies to unify spatial coordinates across slices for k-NN graph construction, each optimized for a given integration scenario. Once the k-NN graph is built, MaskGraphene employs a graph attention autoencoder architecture^23, 24^ to refine low-dimensional latent embeddings by jointly optimizing a masked self-supervised loss (*L*_*masked*_ + *L*_*latent*_ in Fig.1)^25–27^ and a triplet loss. The triplet loss is iteratively minimized by optimizing triplets in the latent space, which act as indirect “soft-links”, further strengthening interslice connections.

**Fig 1.**
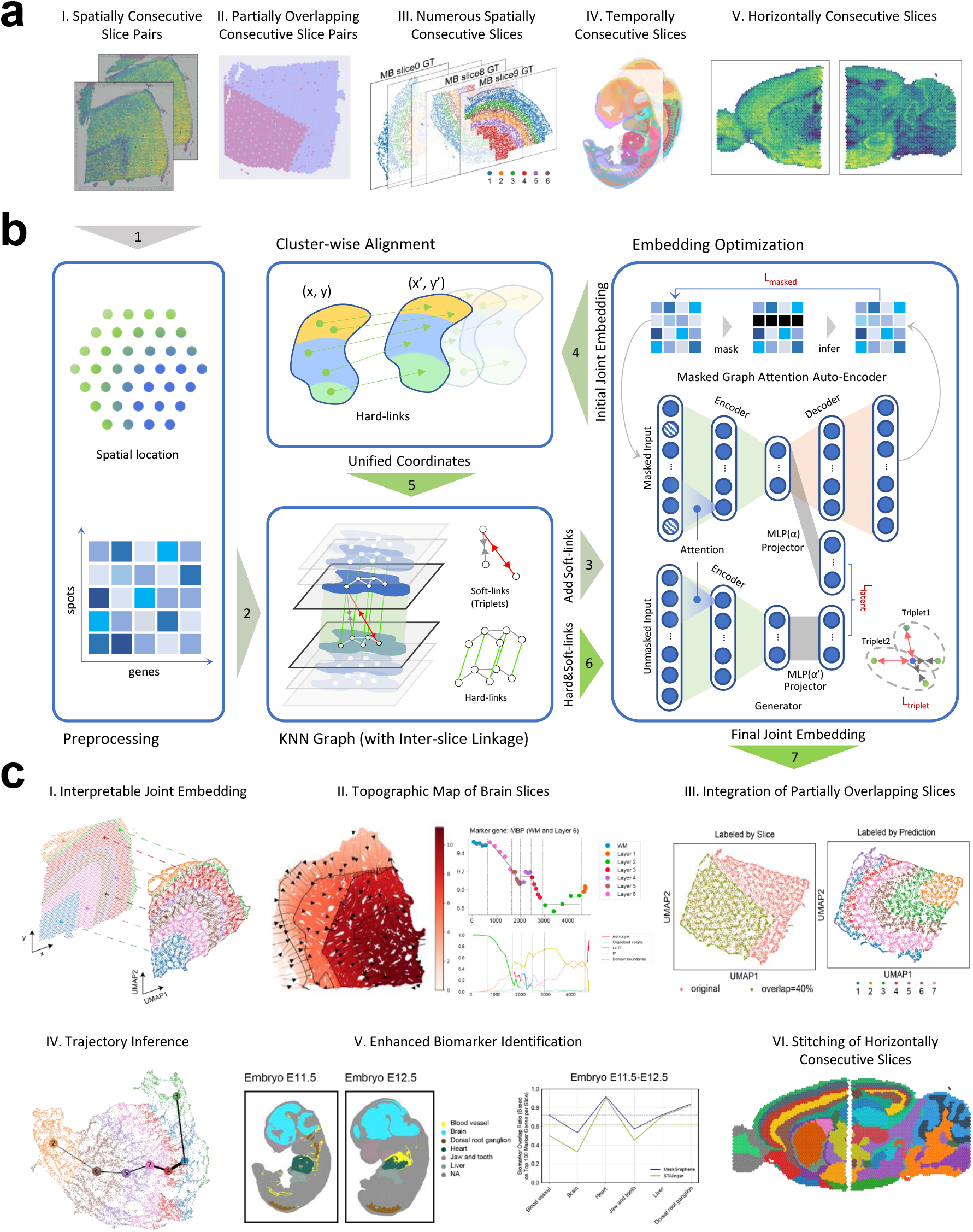
MaskGraphene workflow. **(a)** Illustration of spatial transcriptomics data integration scenarios addressed by MaskGraphene, including spatially consecutive slice pairs (I), simulated partially overlapping consecutive slice pairs (II), numerious spatially consecutive slices (III), temporally consecutive slices (IV), and horizontally consecutive slices (V). **(b)** Workflow of MaskGraphene: The preprocessing step organizes spatial coordinates and gene expression data. Interslice linkage is established through the construction of “hard-links” via cluster-wise alignment and “soft-links” using contrastive learning with triplets. Embedding optimization leverages a masked graph autoencoder to generate batch-corrected joint embeddings by optimizing masked self-supervised loss (*L*_*masked*_ + *L*_*latent*_) and triplet loss. **(c)** Applications and evaluations: (I) Interpretable joint embedding captures the original geometric structure. (II) Topographic map of brain slices with isodepth analysis reveals gene expression gradients across cortical layers. (III) Validation with simulated data demonstrates robust integration of partially overlapping slices. (IV) Trajectory inference reveals linearly connected developmental trends. (V) Alignment and integration of embryonic tissue structures enhance biomarker identification. (VI) Stitching of horizontally consecutive slices reconstructs spatially coherent regions.

Notably, during the initial clustering phase of the cluster-wise alignment method, the masked selfsupervised loss and triplet loss (“soft-links”) are integrated within the graph attention autoencoder to learn initial joint embeddings for clustering. Detailed methods for MaskGraphene are provided in the Methods section.

We benchmarked MaskGraphene against seven integration methods under various experimental scenarios: BASS, DeepST, PRECAST, GraphST, SPIRAL, STAligner, and SpaDo. The details for each tool are provided in Supplementary Table 1.

### Enhanced alignment and mapping accuracy achieved through joint embeddings in MaskGraphene

To assess whether MaskGraphene generates superior joint latent embeddings for integrating consecutive slices compared existing integration methods such as DeepST, STAligner, GraphST, PRECAST, SPIRAL, and SpaDo, we conducted two evaluation experiments involving nine pairs of human dorsolateral prefrontal cortex (DLPFC) slices and four pairs of mouse hypothalamus (MHypo) slices. The DLPFC dataset, generated using 10x Visium, includes 12 human sections with annotations for cortical layers 1–6 and white matter (WM) from three samples.^28^ Each sample has four consecutive slices (A, B, C, D), with AB and CD being 10 *µ*m apart and BC separated by 300 *µ*m. The MHypo dataset by MERFISH consists of five consecutive slices,^17^ labeled Bregma - 0.04mm, -0.09mm, -0.14mm, -0.19mm, and -0.24mm, each with detailed cell annotations. Additional details about these datasets can be found in the Methods section.

The first evaluation focused on layer-wise alignment accuracy,^11^ based on the critical hypothesis that aligned spots across adjacent consecutive slices are more likely to belong to the same spatial domain or cell type. We utilized joint embeddings learned from all tools, based on Euclidean distance, to align (anchor) spots from the first slice to the corresponding (aligned) spots on the second slice for each slice pair. In Fig.2a, we compared the layer-wise alignment accuracy of seven methods across all nine DLPFC slice pairs. Leveraging the distinctive layered structure of the DLPFC data, this evaluation metric was designed to assess whether anchor and aligned spots belonged to the same layer (layer shift = 0) or to different layers (layer shift = 1 to 6). An effective integration tool was expected to achieve high accuracy for anchor and aligned spots within the same layer, with accuracy progressively decreasing as the layer shift increased. In Fig.2a, we plotted the layer-wise alignment accuracy and ranked the tools in descending order. MaskGraphene consistently exhibited the highest accuracy (for a layer shift of 0) across all nine DLPFC slice pairs, outperforming all other tools. For all tools, the majority of anchor and aligned spots across two consecutive slices were mapped to the same layer. However, distant DLPFC slice pairs (300*µ*m apart), such as DLPFC 151508-151509, presented challenges for all methods except MaskGraphene. In these cases, layer-wise alignment accuracy (for a layer shift of 0) significantly decreased compared to near slice pairs (10*µ*m apart), with a large proportion of anchor spots aligning to the adjacent layers (1 or 2 layers away).

**Fig 2.**
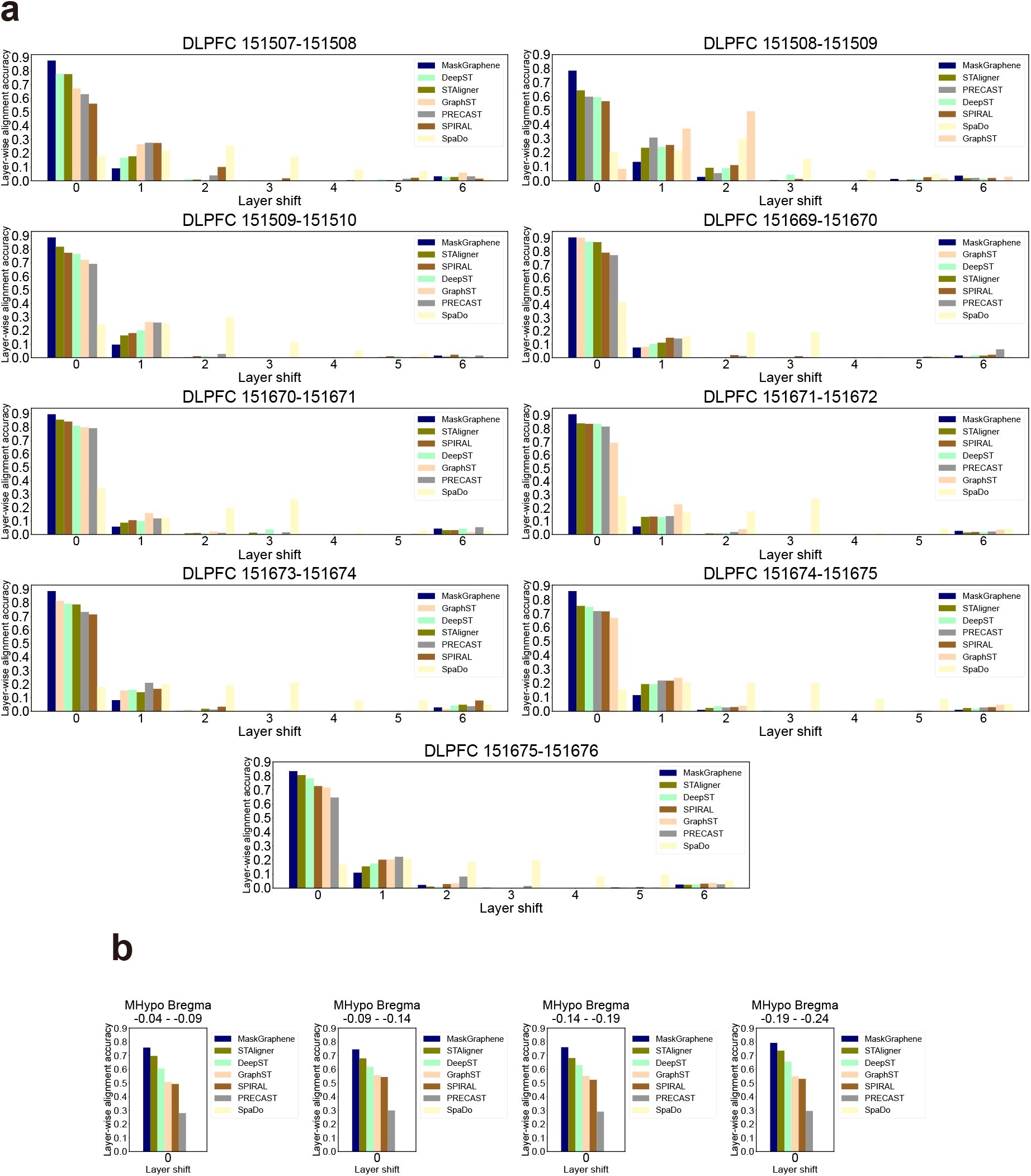
Bar plots for layer-wise alignment accuracy on DLPFC and MHypo datasets. **(a)** Bar plots illustrating the layer-wise alignment accuracy across layer shifts ranging from 0 to 6, comparing seven various methods on nine DLPFC slice pairs. **(b)** Bar plots illustrating the layer-wise alignment accuracy for a layer shift of 0, comparing seven various methods on four MHypo slice pairs. Tools are sorted in descending order based on the accuracy for layer shift of 0 in **(a, b)**.

A similar experiment was performed on four pairs from the MHypo dataset (Fig.2b). However, layerwise alignment accuracy was only assessed for a layer shift of 0, as the hypothalamus lacks a distinct layered structure. MaskGraphene outperformed all other methods, with STAligner and DeepST ranking second and third, respectively. We attribute MaskGraphene’s high alignment accuracy to its use of interslice “hard-links”, which effectively facilitate the connection of similar spots or regions across slices.

To further evaluate the quality of the joint embeddings, we calculated the spot-to-spot matching ratio,^11^ a metric that quantifies how well spots on one slice align with their corresponding spots on the adjacent slice. For the DLPFC 151507–151508 pair (Fig.3a), we annotated ‘anchor’ and ‘aligned’ spots on both slices, using three distinct colors. Based on the ground truth layer labels, these spots were further categorized as aligned (orange), misaligned (blue), and unaligned (green). MaskGraphene (1.27), GraphST (1.38), and PRECAST (1.85) exhibited lower spot-to-spot matching ratios (below 2), indicating superior one-to-one alignment fidelity. In contrast, methods such as SpaDo (4.36), STAligner (2.82), SPRIAL (2.41), and DeepST (2.13) showed higher matching ratios, reflected in an increased number of unaligned spots on the second slice. This is likely due to a tendency to map multiple anchor spots (2-5) to a single aligned spot, resulting in a many-to-one integration pattern. While this may still achieve good layer-wise alignment, it leads to a higher spot-to-spot matching ratio. These trends were consistently observed on other DLPFC pairs of different samples by all tools, as shown in Supplementary Fig.1. Averaging the spot-to-spot ratio across all nine DLPFC pairs (Fig.3c) showed that MaskGraphene (1.31) consistently achieved the lowest ratio, indicating the best performance, followed by GraphST (1.38), PRECAST (1.85), and DeepST (2.13). In contrast, tools such as SPIRAL (2.41), STAligner (2.78), and SpaDo (3.30) exhibited higher ratios. Moreover, across all nine pairs, misaligned spots on the first slice were observed to aggregate along layer boundaries as thin, tightly clustered layers in MaskGraphene and GraphST. Conversely, the other tools showed a more dispersed pattern of misaligned spots within the layers. These results indicate that while better performing tools primarily struggle with aligning spots near domain boundaries, worse performing tools misalign spots even within individual domains.

**Fig 3.**
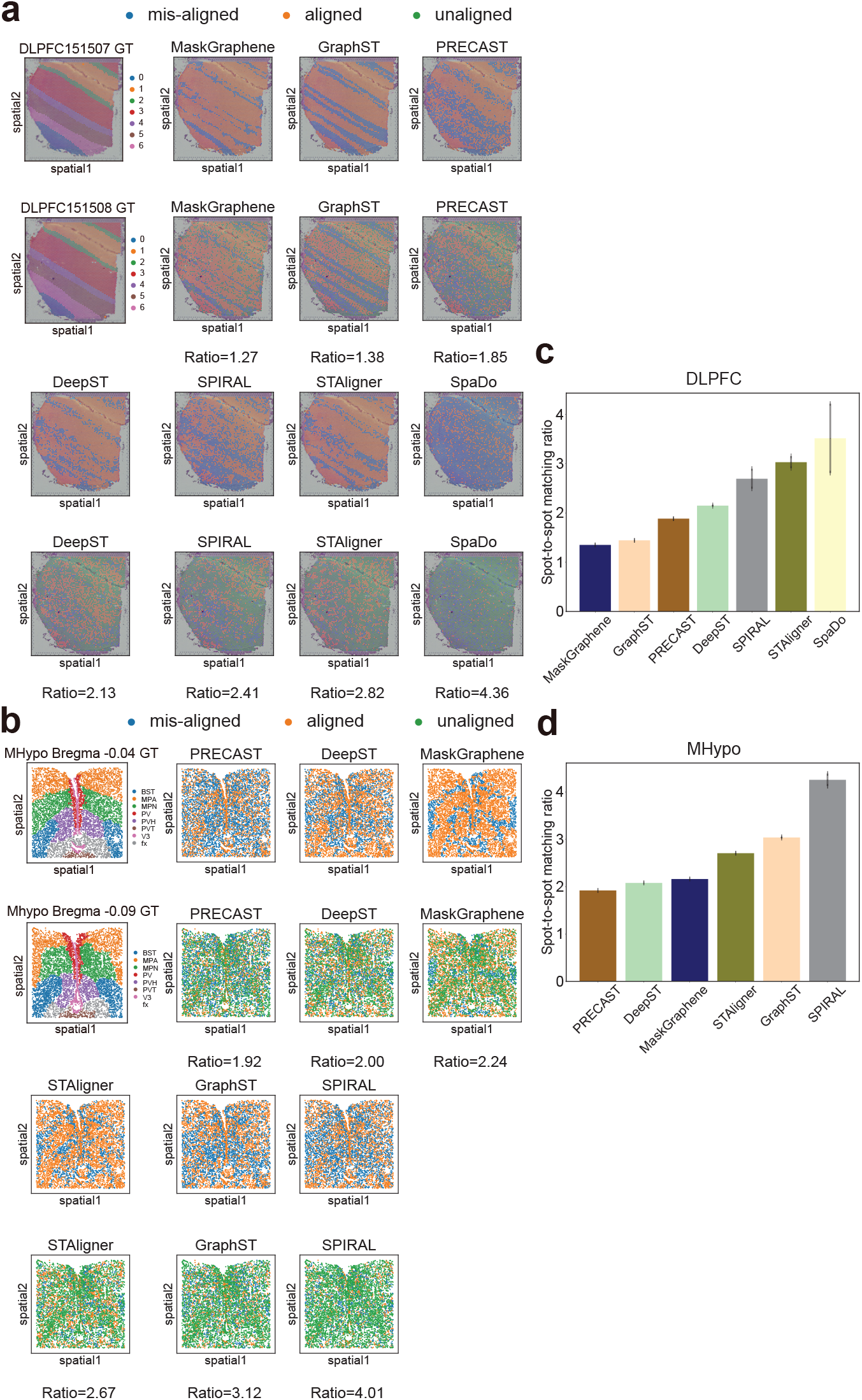
Visualization plots for alignment-misalignment-unalignment and spot-to-spot mapping ratio on DLPFC and MHypo datasets. **(a-b)** Visualization plots displaying aligned spots, misaligned spots, and unaligned spots during the alignment process. The anchor spot from the first (top) slice is aligned to the corresponding spots on the second (bottom) slice for DLPFC 151507-151508 pair **(a)** and MHypo Bregma -0.04 - -0.09 pair **(b)**. The first slice pair illustrating ground truth (GT) annotations. The values below each plot indicate the spot-to-spot matching ratio. **(c-d)** Bar plots representing the average spot-to-spot mapping ratio of each tool on two datasets: DLPFC **(c)** and MHypo **(d)**.

For the MERFISH MHypo dataset, unlike the DLPFC dataset, the transcriptomic spots exhibit an irregular spatial distribution, and the slice pairs are separated by a greater distance (0.05 mm apart). Similar to the decreased layer-wise alignment accuracy (Fig.2b vs. Fig.2a), the performance of all methods either declined or remained poor in terms of spot-to-spot matching ratios compared to the DLPFC dataset, as shown in Fig.3b and d and Supplementary Fig.2. Specifically, PRECAST (1.92), DeepST (2.00), and MaskGraphene (2.24) exhibited relatively lower ratios compared to STAligner (2.67), GraphST (3.12), and SPRIAL (4.01). The average ratio across all four pairs in Fig.3d reflected a similar trend. Although PRECAST and DeepST achieved slightly lower matching ratio than MaskGraphene, their misaligned spots were scattered within the spatial domains, consistent with their pattern observed in the DLPFC dataset (Fig.3b and Supplementary Fig.2). In contrast, MaskGraphene exhibited thin, tightly aggregated lines of misaligned spots near the domain boundaries.

In summary, MaskGraphene demonstrates the best alignment and mapping performance, achieved through its joint embeddings. For deep learning-based methods, it is common for spots in low-dimensional space to lose some geometric information from the original gene expression and spatial coordinate profile during optimization. As a result, these tools tend to exhibit poorer spot-to-spot alignment performance, but achieve better layer-wise alignment accuracy. By introducing “hard-links” to strengthen interslice connections, MaskGraphene imposes additional constraints on the optimization process, enabling its joint embeddings to better preserve the original geometric structure and improve spot-to-spot alignment.

### MaskGraphene enhances integration with interpretable joint embedding to preserve the original geometric structure and reconstruct spatial trajectory

In the previous section, we evaluated the quality of joint embeddings through pairwise evaluations of slice alignment and spot mapping. Expanding on this analysis, we further evaluated the integration quality of the “batch-corrected” joint embeddings generated by MaskGraphene using uniform manifold approximation (UMAP) visualizations for both DLPFC and MHypo slices. These evaluations were conducted in both pairwise and multi-slice scenarios for each dataset, comparing MaskGraphene with benchmarked methods. Each UMAP visualization is color coded based on ground truth (GT) annotations, predicted domains, and the source of origin, shown in left, middle, and right panels, respectively, for each tool (Fig.4a-b).

**Fig 4.**
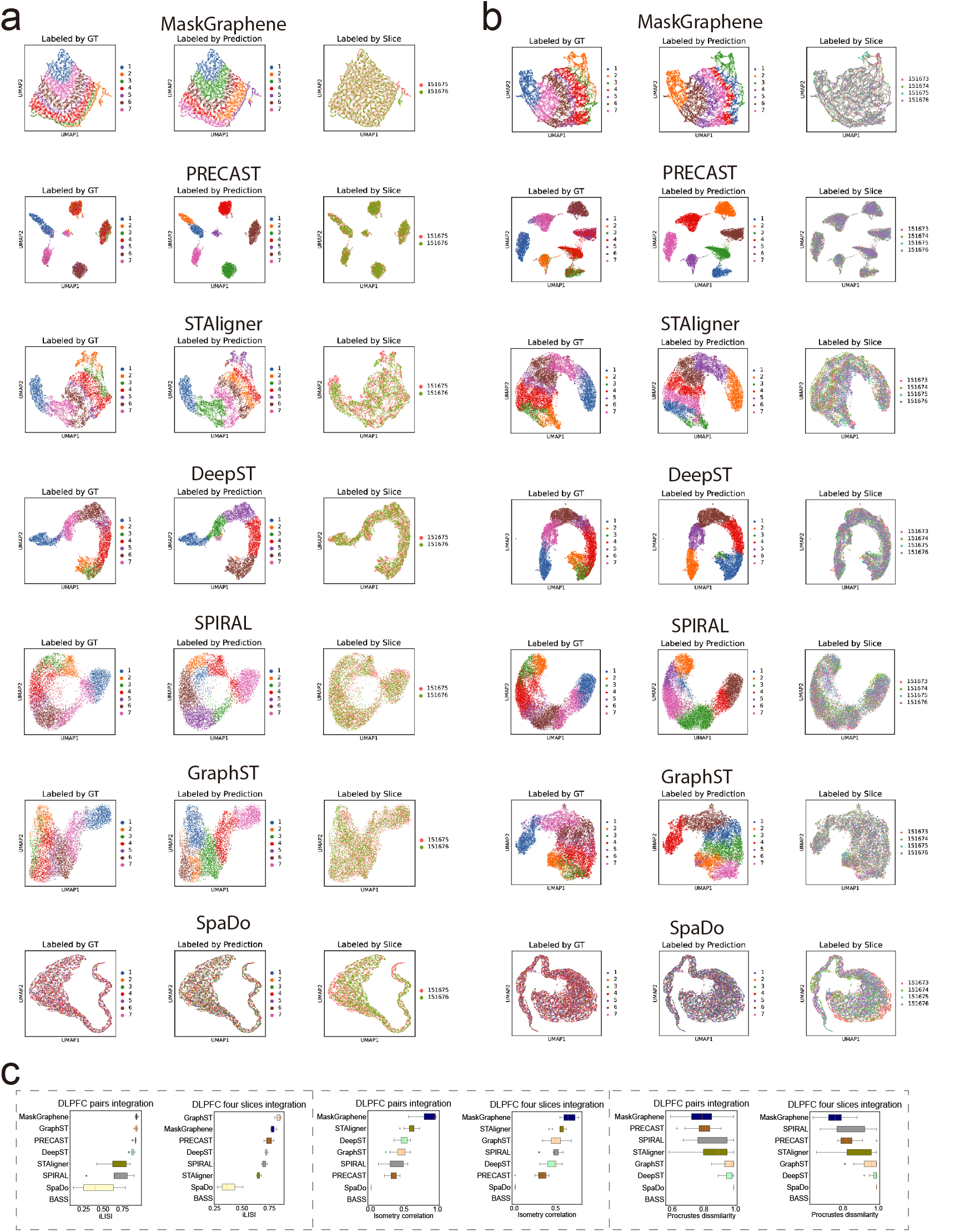
UMAP plots of low dimensional joint embedding and box plots for iLISI, Isometry correlation, and Procrustes dissimilarity on the DLPFC dataset. **(a)** UMAP visualizations of joint embeddings generated by different methods for the DLPFC 151675-151676 pair-wise integration. Spots are colored by ground truth (GT) labels, predicted domains, and slice identity. Each row corresponds to a method: MaskGraphene, PRE-CAST, STAligner, DeepST, SPIRAL, GraphST, and SpaDo. **(b)** UMAP visualization on DLPFC four-slice integration (151673-151674-151675-151676). **(c)** Left Panels: Box plots representing iLISI scores for different integration methods across all DLPFC pair-wise and four-slice integrations. Middle Panels: Box plots representing isometry correlation scores across all DLPFC pair-wise and four-slice integrations. Right Panels: Box plots representing Procrustes dissimilarity scores across all DLPFC pair-wise and four-slice integrations.

Starting with the DLPFC 151675-151676 pair, UMAP plots for all tools showed that spots from the two slices were evenly mixed (Fig.4a, right panels), and their predicted domain clusters were generally well segregated (Fig.4a, middle panels). However, the concordance with the ground truth varied across tools (Fig.4a, middle panels vs. left panels). Specifically, PRECAST tended to generate embeddings with highly separated clusters, often sacrificing geometric information. This led to predicted clusters that encompassed spots from different domains, a pattern that did not align well with the ground truth. SpaDo displayed the worst integration pattern. In contrast, tools such as MaskGraphene, STAligner, DeepST, SPIRAL, and GraphST preserved the hierarchical connections of the seven layers in the latent embedding space to varying degrees. These tools occasionally predicted spatial domains that included a few spots from nearby domains or split a single spatial domain into two adjacent ones. Notably, MaskGraphene demonstrated a unique ability to recover the entire slice shape, layer-wise patterns, and spatial relationships in the UMAP visualization.

Overall, MaskGraphene produced the most visually coherent UMAP results among all tools, effectively mitigating batch effects and demonstrating superior integration quality. For the four-slice integration scenario (DLPFC 151673-151674-151675-151676) shown in Fig.4b), similar UMAP visualizations and trends were observed across all tools. MaskGraphene continued to produce the most visually coherent UMAP results among all methods, although the overall UMAP outline was slightly deformed compared to the pairwise integration case. Similar UMAP visualization patterns were also observed in other pairwise and multi-slice scenarios, as demonstrated in Supplementary Fig.3.

To qualitatively assess the batch effect removal and geometry preservation in the embeddings, we employed three metrics, as shown in Fig.4c. Detailed descriptions of these metrics are included in the Methods section. Batch effect removal across different methods was quantified using iLISI, where MaskGraphene achieved the highest scores in pairwise integration scenario and the second-highest in four-slice integration scenario, reflecting its ability to maintain well-mixed embedding across slices. Furthermore, MaskGraphene achieved the best performance in both pairwise and four-slice integration scenarios based on two geometry-preservation metrics: Isometry correlation and Procrustes dissimilarity. A higher Isometry correlation score indicates better geometry preservation, while a lower Procrustes dissimilarity score reflects superior geometry preservation. These findings align with the advanced alignment and mapping accuracy achieved by joint embeddings of MaskGraphene, further validating its capability to recover geometric shapes, spatial patterns, and layerwise structures across various DLPFC slice integration scenarios.

We then performed similar UMAP and qualitative analyses on the MHypo dataset for both pairwise and five-slice integration scenarios. As shown in Fig.5a, MaskGraphene demonstrated exceptional integration with batch correction compared to the other six tools. Its predicted domain clusters were well segregated and highly concordant with the ground truth (Fig.5a, middle panels vs. left panels). The joint embeddings generated by MaskGraphene retained some degree of the original geometric information, although this effect was less pronounced compared to the DLPFC data. In contrast, the other six tools displayed significantly poorer integration performance relative to the ground truth, particularly in the four-slice integration scenarios (Fig.5b). To further evaluate the joint embeddings using the three metrics applied to the DLFPC data, we found that MaskGraphene achieved the best scores across all integration scenarios on the MHypo dataset, with the exception of the iLISI score in the pairwise integration scenario.

**Fig 5.**
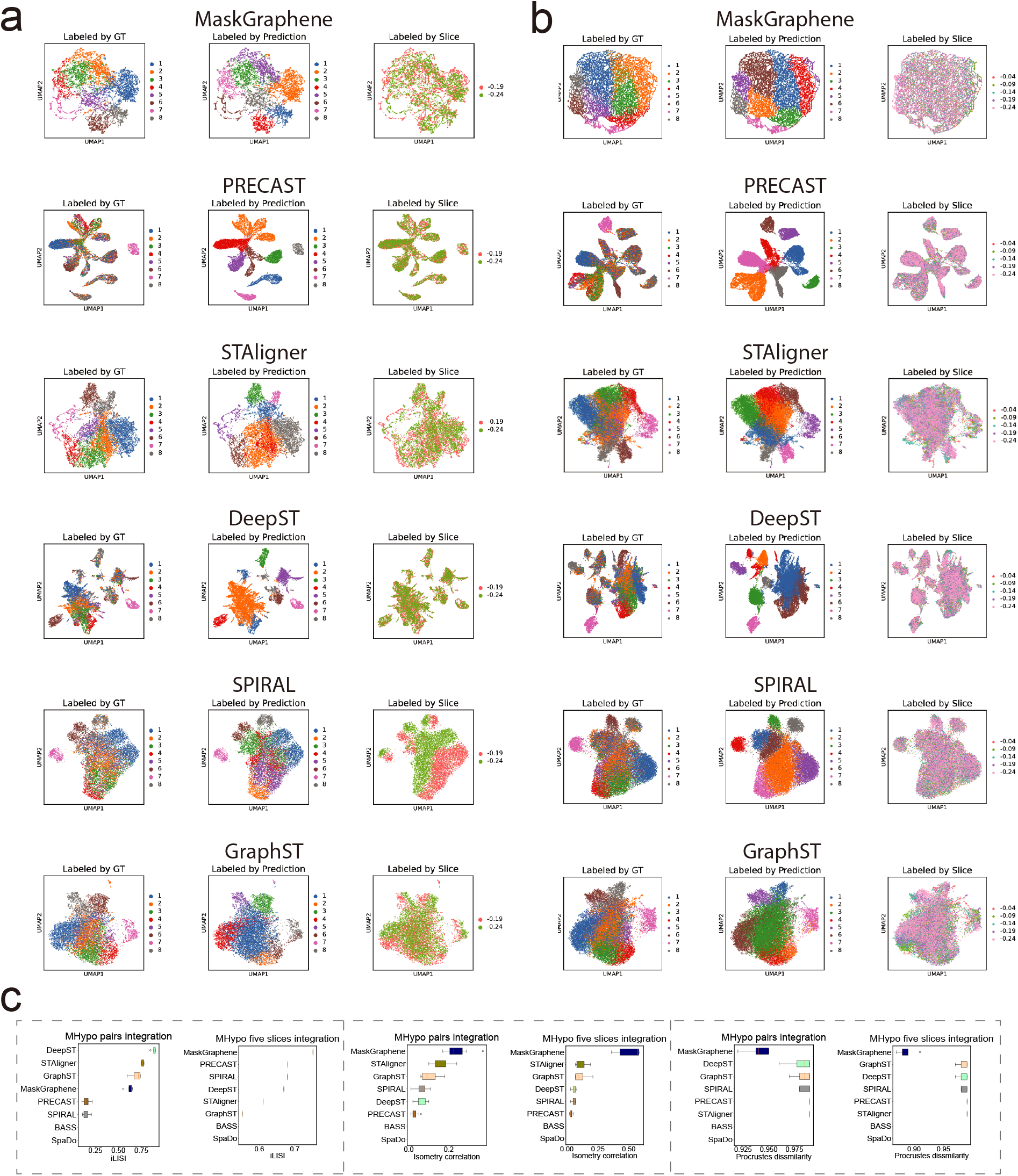
UMAP plots of low dimensional joint embedding and box plots for iLISI, Isometry correlation, and Procrustes dissimilarity on the MHypo dataset. **(a)** UMAP visualizations of joint embeddings generated by different methods for the MHypo Bregma -0.19 - -0.24 pair-wise integration. Spots are colored by ground truth (GT) labels, predicted domains, and slice identity. Each row corresponds to a method: MaskGraphene, PRECAST, STAligner, DeepST, SPIRAL, and GraphST. **(b)** UMAP visualization on MHypo five-slice integration (Bregma -0.04 - -0.24). **(c)** Left Panels: Box plots representing iLISI scores for different integration methods across all MHypo pair-wise and five-slice integrations. Middle Panels: Box plots representing isometry correlation scores across all MHypo pair-wise and five-slice integrations. Right Panels: Box plots representing Procrustes dissimilarity scores across all MHypo pair-wise and five-slice integrations. BASS can not generate embeddings for UMAP visualizations, while SpaDo is incompatible with the MHypo dataset.

MaskGraphene effectively revealed spatial domains and boundaries by projecting joint embeddings onto a two-dimensional space using UMAP visualizations on the DLPFC dataset. We thus further inferred spatial trajectories with the trajectory inference tool PAGA to benchmark the quality of the joint embeddings. In Fig.6, we present PAGA graphs illustrating connectivity patterns among spatial domains after four-slice integration using MaskGraphene, STAligner, and GraphST. We selected STAligner, and GraphST due to their relatively strong integration performance. Each set of two figure panels displays UMAP visualizations alongside PAGA graphs, with slices labeled by predicted clusters (Fig.6). Across all three different four-slice integration scenarios, the PAGA graphs and UMAP plots generated by MaskGraphene’s embeddings showed that clusters corresponding to each layer were distributed accurately and exhibited the most consistent spatial trajectory from layer 1 to layer 6 and white matter (WM), indicating a linearly connected developmental trend. In contrast, STAligner demonstrated moderate connectivity, with some mixed connections in the middle layers, but struggled to maintain consistency. GraphST produced fragmented PAGA graphs, highlighting its limitations in preserving spatial domain continuity.

**Fig 6.**
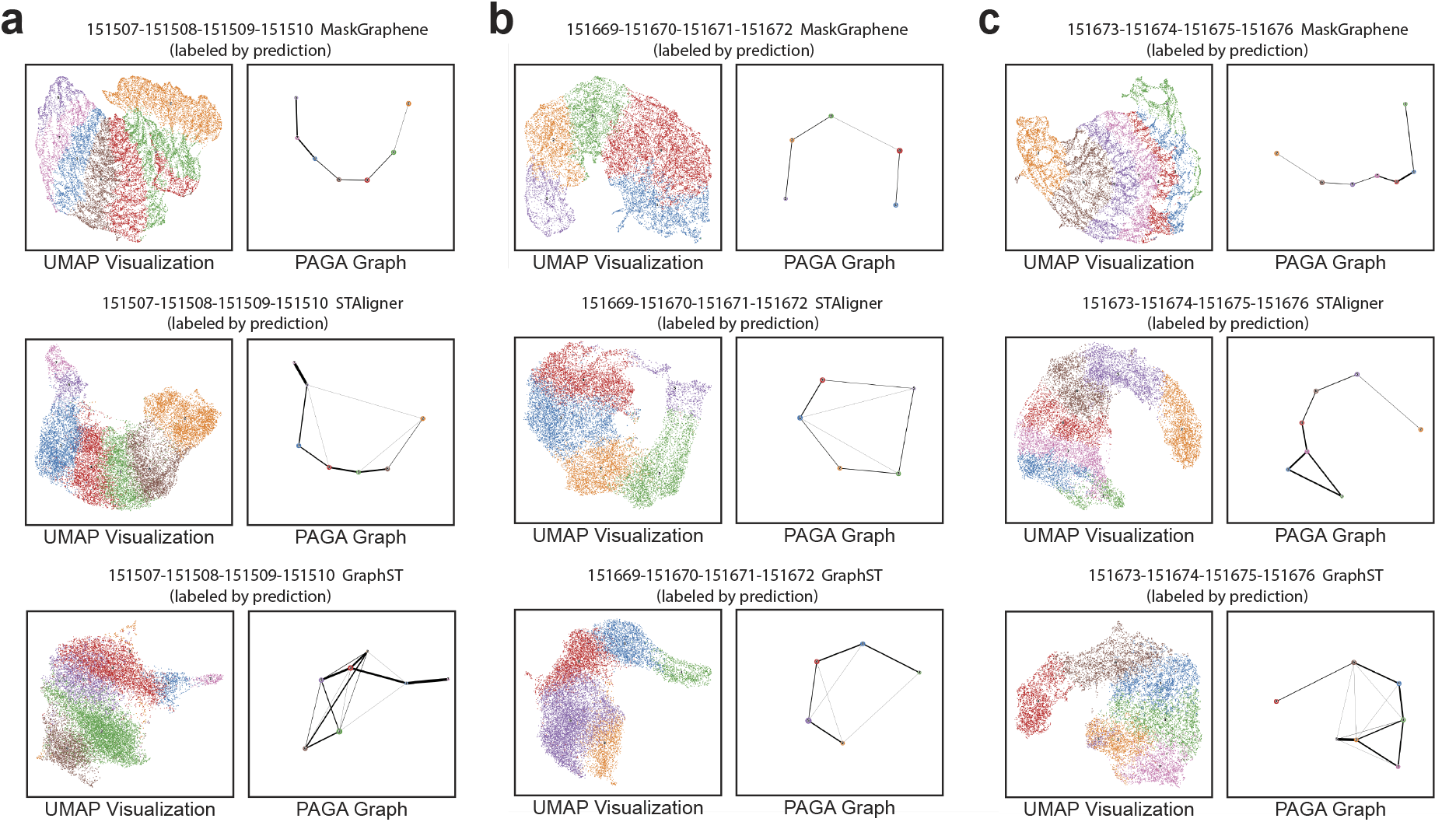
Visualization of UMAP paired with PAGA graphs on the DLPFC dataset. **(a-c)** Each panel shows UMAP visualizations paired with PAGA graphs, illustrating spatial trajectory results for three distinct DLPFC four-slice integration: (151507-151508-151509-151510) **(a)**, (151669-151670-151671-151672) **(b)**, and (151673-151674-151675-151676) **(c)**. Spots are colored according to predicted domains. Each row corresponds to a method: MaskGraphene, STAligner, and GraphST.

In summary, the results suggest that MaskGraphene’s batch-corrected joint embeddings, achieved through pair-wise and multi-slice integration, effectively preserve the original geometric structure and accurately reconstruct spatial trajectory. Although other methods demonstrate competitive performance, they fall short of achieving the same level of robustness and precision. These findings highlight MaskGraphene’s capability to produce interpretable joint embedding in spatial transcriptomics, facilitating robust integration and efficient batch effect correction.

### MaskGraphene improves biomarker analysis and uncovers the topography of brain slices through interpretable joint embeddings

Next, we sought to evaluate MaskGraphene’s performance in improving biomarker identification and demonstrate its ability to derive a topographic map of brain slices through joint embeddings, revealing gradients of neuronal differentiation and activity. To assess whether integration improves biomarker identification, we compared layer marker genes identified from individual slices based on ground truth labels with biomarkers identified from each integrated layer across four slices. First, we matched MaskGraphene’s predicted domains with ground truth labels to annotate the layers (Fig.7a) and extracted spots corresponding to each layer from all four slices. Biomarkers for each integrated layer were then identified using Scanpy.^29^ Next, to generate ground truth layer marker genes for comparison, we used Scanpy to identify biomarkers for each layer in individual slices based on ground truth labels, selecting the top 50 or 100 genes per layer per slice. The union of these top genes, denoted as *N*, across all four slices was compiled as the final set of ground truth layer marker genes for validation. Finally, we calculated the overlap ratio between the top *N* biomarkers identified for each integrated layer and the *N* ground truth layer marker genes. MaskGraphene’s performance was benchmarked against STAligner and GraphST. As shown in Fig.7a, MaskGraphene’s predictions closely matched the ground truth, accurately capturing the spatial organization of these brain regions. Furthermore, MaskGraphene achieved higher overlap ratio across nearly all layers when compared to the ground truth marker genes, defined by selecting the top 50/100 layer marker genes (Figure 7b). This performance exceeded that of STAligner and GraphST, particularly in layers 1 and 3, where MaskGraphene exhibited significantly higher overlap ratios. These results highlight MaskGraphene’s enhanced accuracy in identifying and localizing these domains after integration, thereby enhancing biomarker identification.

**Fig 7.**
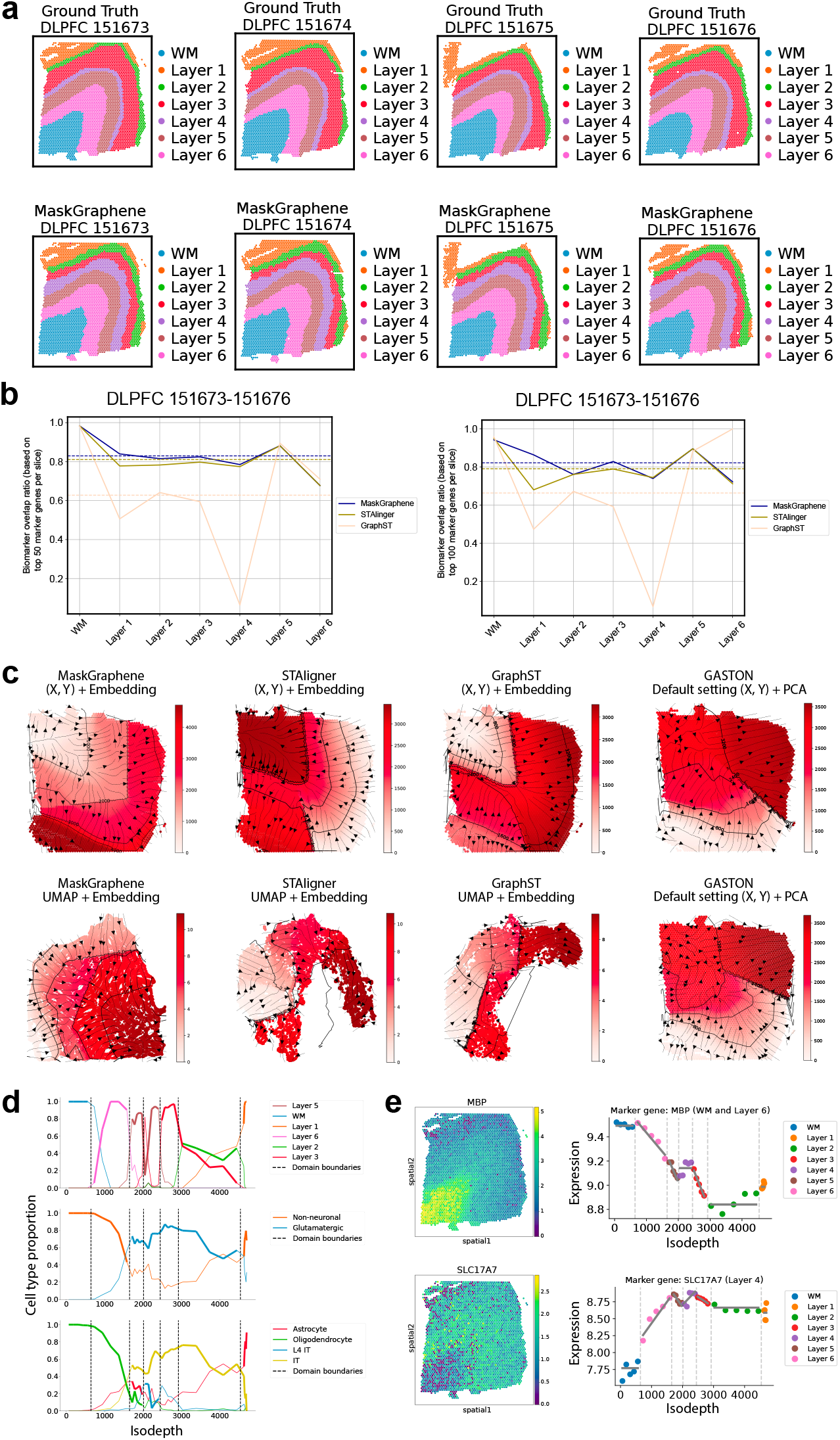
Biomarker and topography analysis after brain slice integration. **(a)** Spatial visualizations of joint domain identification for four DLPFC slices (151673, 151674, 151675, and 151676) based on ground truth (top panels) and MaskGraphene predictions after integration (bottom panels). Colored annotations indicate white matter (WM) and six cortical layers (1–6) across two conditions. **(b)** Biomarker overlap ratio curves across WM and cortical layers for three different tools, comparing the top *N* biomarkers identified for each integrated layer after four-slice integration with the *N* ground truth marker genes. Ground truth markers are defined as the union of the top 50 (left panel) or top 100 (right panel) layer marker genes across four slices. **(c)** Top panels: Topographical maps generated using GASTON with four settings: original X,Y coordinates combined with joint embeddings after DLPFC 151673-151674 pair-wise integration from MaskGraphene, STAligner, GraphST, and default PCA-derived embeddings from DLPFC slice 151673. Bottom panels: Maps generated with UMAP coordinates combined with embeddings from MaskGraphene, STAligner, GraphST, and default X,Y coordinates with PCA-derived embeddings from slice 151674. **(d)** Plots showing the proportions of cell types as a function of the isodepth, using three different types of annotations: layer-specific cell types (top panel), neuronal types (middle panel), and cell types (bottom panel). **(e)** Layer-specific marker gene analysis for MBP (white matter, layer 6) and SLC17A7 (layer 4). The left panels exhibit heatmaps of gene expression intensity, while the right panels depict the marker gene expression versus the isodepth.

We next employed GASTON^3^ to analyze the topography of brain tissue slices. By integrating PCA-derived spot embeddings with spatial information, GASTON can characterize the topography of individual tissue slices. To assess whether joint embeddings from integration provide improved topographical representations and facilitate downstream gene expression gradient analyses compared to single-slice analysis, we utilized MaskGraphene’s joint embeddings as GASTON’s input. Additionally, to evaluate the extent to which two-dimensional UMAP coordinates preserve the original geometric structure, we replaced the actual X,Y coordinates with UMAP coordinates as input. To benchmark performance, we compared MaskGraphene against established tools such as STAligner and GraphST under various control settings, including GASTON’s default single-slice configuration.

As illustrated in Fig.7c, the learned isodepth contour lines delineate a topographical map of the DLPFC slices, identifying the boundaries between distinct cortical layers. The spatial expression gradients, oriented perpendicular to these cortical layers (constant isodepth contours), indicate the directions of maximum gene expression change. By comparing all topographical maps by different settings, we found that GASTON, leveraging MaskGraphene’s joint embeddings from pairwise integration of DLFPC 151673-151674 pair along with X,Y coordinates, effectively segmented the tissue into several contiguous spatial domains that visually align with the layered structure of the DLPFC. Notably, this approach outperformed single-slice analysis with GASTON using default PCA-derived embeddings and X,Y coordinates. In contrast, when employing joint embeddings from STAligner or GraphST combined with X,Y coordinates, GASTON exhibited poor spatial organization of the DLPFC. Furthermore, when using two-dimensional UMAP coordinates derived from MaskGraphene’s embeddings in place of the original spatial context, we still observed a clear topographical map of the layered geometry of the DLPFC with well-defined continuous gradients. Conversely, GASTON failed to capture the topographical map when using the UMAP derived from STAligner and GraphST as input. These results further validates that MaskGraphene’s joint embeddings effectively capture the geometric structure, enabling robust downstream analyses.

We further compared the spatial domains (cortical layers) to the cell types reported in the original publication for this dataset. These cell types were obtained from the deconvolution module of SPACEL,^30^ which performs cell type deconvolution based on a reference scRNA-seq dataset. To conduct spatial domain-isodepth and cell type-isodepth analyses,^3^ we utilized the topographical map derived from integrating MaskGraphene’s joint embeddings with the original spatial context. Specifically, we computed isodepth coordinates, enabling us to examine continuous variations in cell types within and across the cortical layers of the DLPFC. Our analysis revealed significant variations in cell type proportions along the isodepth axis (Fig.7d, middle and bottom panels), which aligned well with the reported cell type abundances and their functional roles within each cortical layer (Fig.7d, top panel). For example, by aligning and analyzing the three panels together (Fig.7d), we observed that a high abundance of oligodendrocytes across the isodepth range corresponding to white matter (WM) and the deeper cortical layers (layer 5 and 6). This finding is consistent with their critical role in myelinating the large, long-projecting axons of pyramidal neurons that connect the cortex to subcortical structures.^31^ These oligodendrocytes also constitute a significant proportion of the non-neuronal cells in these layers. Furthermore, a sharp transition in cell-type proportions was observed at the isodepth value used as the boundary (second dashed line) between layers 5 and 6, indicating that the learned isodepth and spatial domains effectively separate non-neuronal cells from neurons. Notably, glutamatergic neurons, including IT cells, exhibited a large and nearly constant proportion across the isodepth range corresponding to layers 2–5, consistent with their role in excitatory signaling within the brain and their importance in various cognitive and motor functions.^32^ These findings demonstrate that the topographical map generated using isodepth not only identifies more spatially coherent domains compared to existing methods but also preserves accurate cell type information.

Finally, we explored whether the learned topography could facilitate the identification of biologically meaningful spatial patterns of gene expression. This analysis focused on capturing both continuous variations in expression within or across spatial domains and abrupt discontinuities in gene expression at the boundaries of adjacent spatial domains.^3^ Details of the analysis are described in the Methods section. For example, in the DLPFC, the marker gene MBP (Myelin Basic Protein) is predominantly expressed in oligodendrocytes, the glial cells responsible for forming and maintaining myelin sheaths, essential for efficient nerve signal conduction.^33^ Its expression peaked in the white matter (WM) layer and showed a continuous gradient across layers 5 and 6 along the isodepth axis (Fig.7e, top panels). However, MBP expression exhibited sharp discontinuities at the boundaries of layer 2, reflecting its low abundance in the superficial layers. This gradient pattern aligned with results from the cell type-isodepth analysis. As another example, the layer 4 marker gene SLC17A7, also known as VGLUT1 (vesicular glutamate transporter 1), serves as a key marker for excitatory glutamatergic neurons due to its role in glutamate signaling. This gene showed peak expression in layer 4, indicating a high density of these neurons, despite reduced visibility on the heatmap due to sparse expression values (Fig.7e, bottom panels). This peak extended into layers 3 and 5, reflecting the laminar organization and enrichment of excitatory neurons in these regions.^34^

We performed similar spatial domain-isodepth, cell type-isodepth, and marker gene gradient analyses using joint embeddings generated by STAligner, GraphST, and default PCA-derived embeddings. However, these methods failed to produce biologically meaningful results comparable to those achieved with MaskGraphene (Supplementary Fig.4-6). Additionally, we extended the analyses using joint embeddings from MaskGraphene’s four-slice integration, which yielded results consistent with those observed in pairwise integration (Supplementary Fig.7).

### MaskGraphene successfully captures the partial-overlap slice phenomenon in simulated data

While real datasets provided some insight into alignment accuracy and integration, they lacked precise spot-to-spot alignment ground truth. To thoroughly investigate alignment accuracy and integration, we simulated datasets with a gold standard for different scenarios to demonstrate the robustness of MaskGraphene (Fig.8). In order to explore capturing different overlap geometric structures in the latent space, we conducted simulations using one DLPFC slice as the reference and generated another slice with varying overlap ratios (20%, 40%, 60%, 80%, and 100%) compared to the reference slice (Fig.8b-c). Further details about the data simulation are provided in the Methods section. This allowed us to examine how the joint embeddings by MaskGraphene could accurately represent these variations. The resulting joint embeddings, obtained by integrating the reference slice with each simulated slice, were visualized in the two-dimensional UMAP space. Spots in the visualization were color-coded based on ground truth (GT) annotations, predicted domains, and the source of origin (Fig.8a).

**Fig 8.**
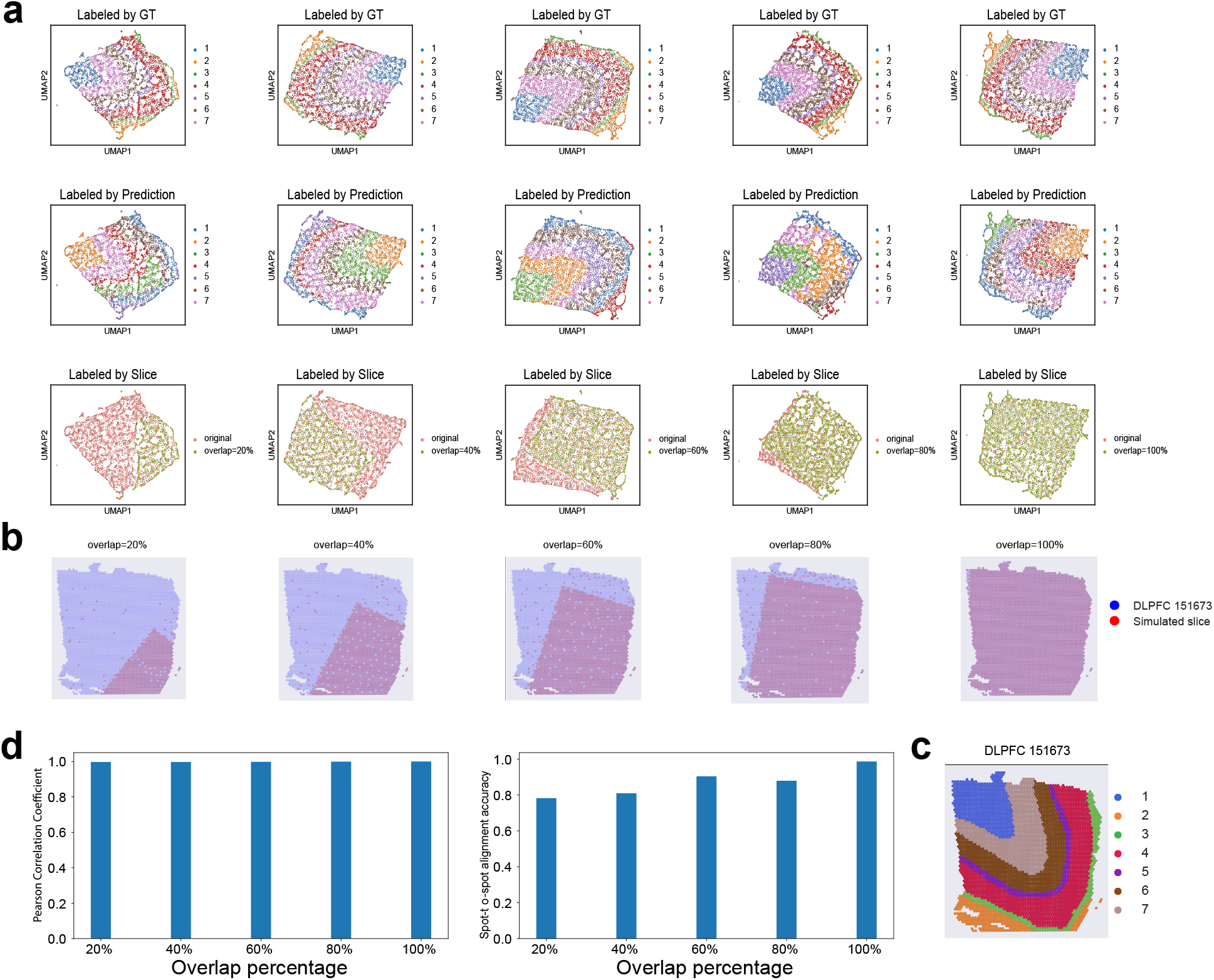
UMAP visualization and validation plots for simulated data integration with varying partial overlap percentages. **(a)** UMAP visualizations labeled by ground truth (GT) labels, predicted domains, and slice identity. Each column represents a specific overlap percentage scenario (20%, 40%, 60%, 80%, and 100%, from left to right). **(b-c)** DLPFC 151673 slice with seven layers, accompanied by simulated consecutive slices showing overlap ratios of 20%, 40%, 60%, 80%, and 100% relative to the reference slice. **(d)** The left panel represents the Pearson correlation between joint embeddings of aligned spots across the reference and simulated slices as the overlap ratio increases. The right panel indicates the spot-to-spot alignment accuracy as the overlap ratio increases.

We noticed that the spots from two different slices were evenly distributed while the predicted cortical layers were clearly separated, showing a strong agreement with the ground truth (Fig.8a, top panels vs. middle panels). The UMAP visualizations preserved the geometric structure of the DLPFC slices, which aligns with our previous observations. Remarkably, the slices labeled by their source effectively depicted the gradually increasing patterns of overlap, showing varying proportions of red spots exclusively associated with the reference slice, as highlighted in the original reference-to-simulated slices (Fig.8a, bottom panels vs. Fig.8b). We computed the Pearson correlation coefficient between joint embeddings of aligned spots based on ground truth and the spot-to-spot alignment accuracy for all five overlap scenarios. As depicted in Fig.8d, the Pearson correlation reached 1.0 for all scenarios, indicating robust embeddings for alignment and integration. The spot-to-spot alignment accuracy achieved 100% when two slices had 100% overlap, and this accuracy slightly decreased as the overlap ratio decreased. This analysis demonstrated that MaskGraphene is capable of integrating partial overlap slices while preserving the underlying geometric structure of the overlaps.

### MaskGraphene enhances domain identification through joint embedding

By integrating data from multiple ST slices, we can estimate joint embeddings of expressions that represent variations between cell or domain types across slices. This approach has the potential to enhance the detection of spatial domains or cell types. To further quantitatively compare the effectiveness of MaskGraphene in capturing spatial domains via joint embeddings, we utilized joint embeddings from pairwise and multi-slice integration in the MHypo and DLPFC datasets to perform clustering using the clustering method mclust.^35^ Subsequently, we calculated the Adjusted Rand Index (ARI) as an evaluation metric to compare the clustering results of MaskGraphene with the ground truth in each slice, with higher ARI scores indicating better domain identification. We benchmarked MaskGraphene against all seven integration methods. BASS was included in this analysis as it provides clustering labels after integration rather than joint embeddings.^11, 17^

In Fig.9a, we plotted the visualization of domain identification with ARI score on each slice for the DLPFC four-slice integration (151673-151674-151675-151676). MaskGraphene demonstrated a distinct separation of the seven-layered regions and achieved the highest clustering accuracy across all four slices. In contrast, most other tools struggled to reveal the expected layer pattern consistently in all slices. Their clustering results often displayed chaotic cluster boundaries and numerous outliers within each cluster, which compromised both the overall clustering accuracy and the visual clarity of the results. Additional visualizations of domain identification with ARI scores for other integration scenarios are provided in Supplementary Fig.8-11. To demonstrate the robustness of clustering performance, we plotted ARI scores across all pair-wise and four-slice integration scenarios using box plots. As shown in Fig.9b, MaskGraphene achieved the best overall clustering performance using joint embeddings from both pairwise and four-slice integration, followed by STAligner and PRECAST.

**Fig 9.**
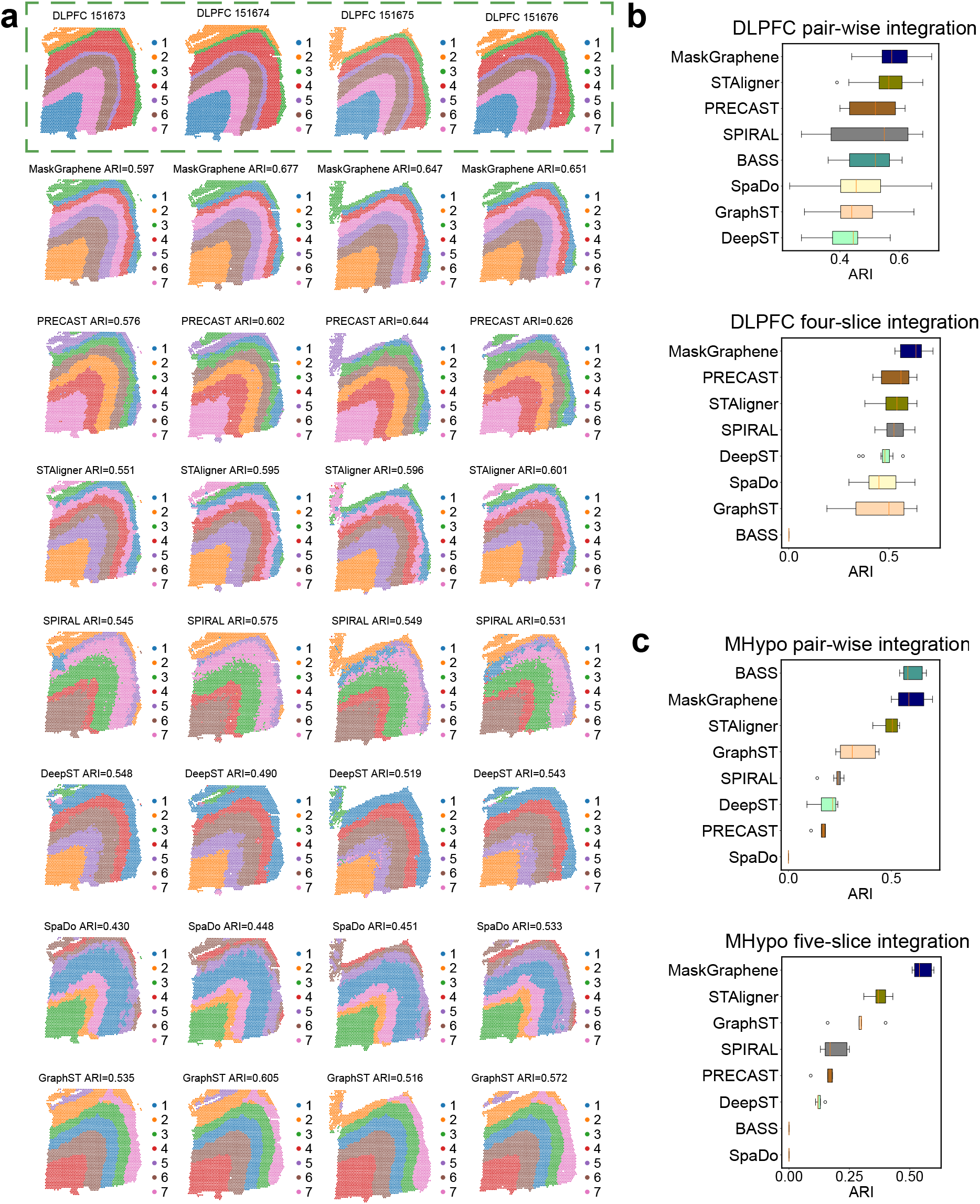
Spatial visualization of joint domain identification and ARI boxplots. **(a)** Spatial domain identification visualization with ARI value for four DLPFC slices (151673, 151674, 151675, and 151676) after integration using seven integration methods, including MaskGraphene, PRECAST, STAligner, SPIRAL, DeepST, SpaDo, and GraphST. The first row, outlined by a green dashed box, represents the ground truth labels. **(b)** Box plots showing ARI scores for clustering results after all DLPFC pair-wise integration (top panel) and four-slice integration (bottom panel) across all methods. Tools are sorted by average ARI. **(c)** Box plot showing ARI scores for all MHypo pair-wise integration (top panel) and the five-slice integration (bottom panel). BASS is not suitable for multi-slice integration, while SpaDo is incompatible with the MHypo dataset.

For the MHypo dataset, as shown in Fig.9c, BASS and MaskGraphene achieved the highest ARI scores, highlighting their superior performance in domain identification after pair-wise integration. In the five-slice integration scenario, MaskGraphene achieved the highest ARI score, followed by STAligner and GraphST, which showed moderate performance. Additional visualizations of spatial domain identification with ARI scores for this dataset are available in Supplementary Fig.12-13.

As outlined in the Methods section, MaskGraphene employs either coordinate replacement or coordinate transformation for multi-slice integration (involving more than two slices) to unify the coordinate system when constructing the k-NN graph. For the DLPFC and MHypo datasets, we primarily demonstrated results obtained using MaskGraphene with coordinate replacement (the default method). Additionally, results generated using MaskGraphene with coordinate transformation are presented in Supplementary Fig.14-15. These results demonstrate that MaskGraphene (coordinate transformation) performs comparably to MaskGraphene (coordinate replacement) across analyses such as UMAP visualization, spatial trajectory, biomarker identification, and spatial domain identification.

### MaskGraphene facilitates the integration of numerous adjacent consecutive ST slices

In this section, we evaluated MaskGraphene’s ability to integrate a substantial number of adjacent consecutive ST slices for large-scale analysis. Specifically, we utilized ten consecutive tissue slices from the primary motor cortex of a mouse embryo (MB) MER-FISH dataset^30, 36^ to perform integration and generate joint embeddings for clustering performance and trajectory inference analysis. Region annotations, including the six layers (L1-L6) and white matter (WM), are provided for each slice for benchmarking. All the other methods, except for STAligner, encountered GPU memory constraints or other issues that prevented them from completing this analysis. In Fig.10a-d, we present the visualization of domain identification, along with the ARI scores for each slice in the MB ten-slice integration. Both MaskGraphene (coordinate replacement) and MaskGraphene (coordinate transformation) consistently outperformed STAligner across nearly all slices, accurately identifying distinct cortical layers with well-defined boundaries as indicated by the ground truth. The high ARI scores achieved by MaskGraphene across all slices (ranging from 0.415 to 0.662) demonstrate its robust and reliable clustering performance, even as the number of integrated slices increased. Overall, MaskGraphene (coordinate transformation) achieved better clustering performance than MaskGraphene (coordinate replacement). In contrast, STAligner exhibited lower ARI scores (ranging from 0.336 to 0.448) and less consistent clustering patterns across all slices. This disparity became more pronounced with an increasing number of slices, indicating that STAligner may struggle to integrate large ST datasets while maintaining clustering accuracy.

**Fig 10.**
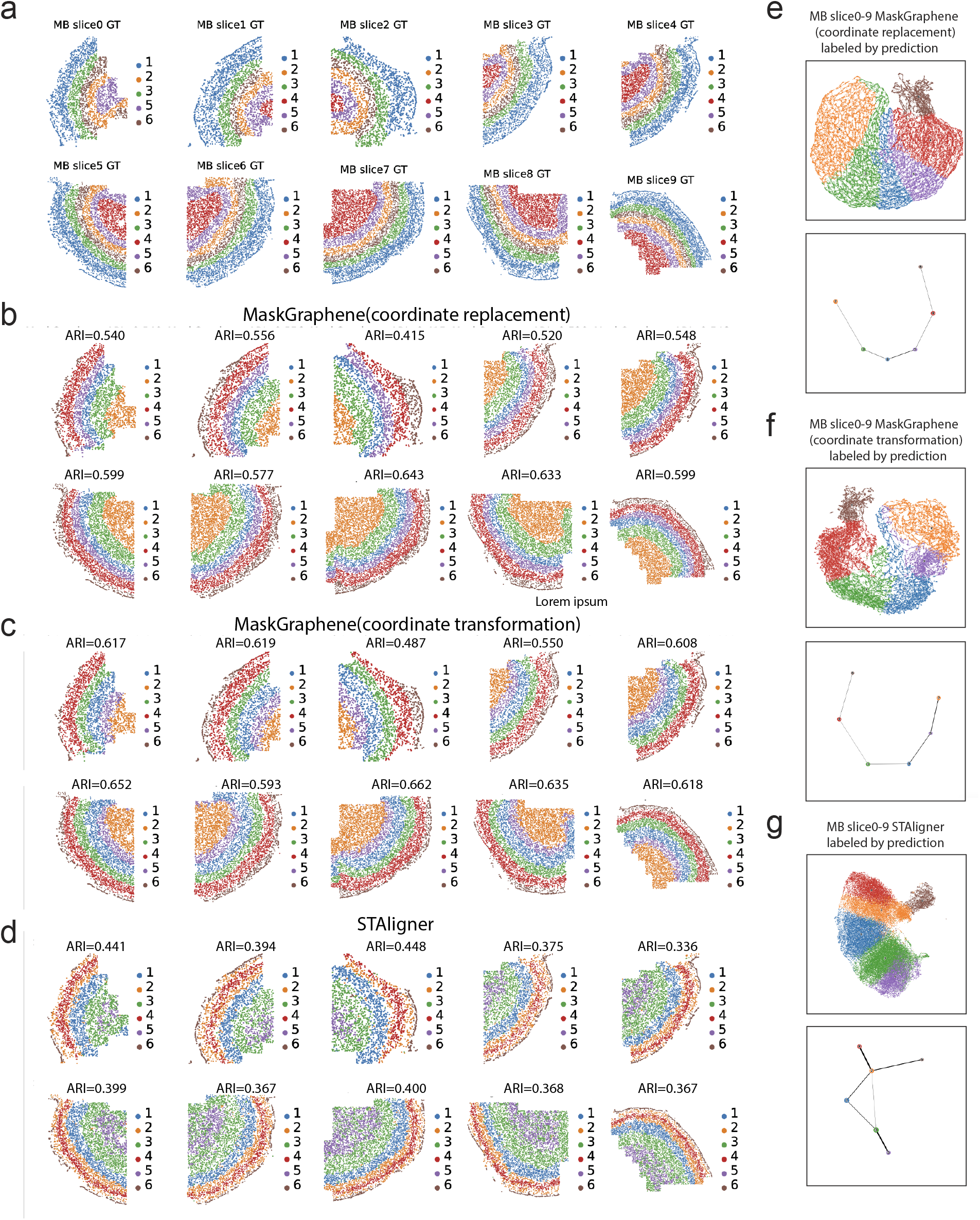
Spatial visualizations of joint domain identification and UMAP paired with PAGA plots for the integration of the MERFISH mouse brain (MB) dataset. **(a)** Spatial domain identification visualization for MB slices (0 - 10) by MaskGraphene and STAligner. Each pair of rows corresponds to a different method or ground truth (GT). **(b)** UMAP visualizations paired with PAGA graphs to assess spatial trajectory results, with UMAP spots colored by predicted domains or GT labels.

The UMAP visualizations (Fig.10e-f, top panels and Supplementary Fig.16) revealed well-distributed clusters corresponding to each layer, displaying a clear hierarchical layer structure with spots from all slices evenly mixed. These results indicate that MaskGraphene effectively harmonized data across numerous slices, successfully mitigating potential batch effects and interslice variability that might otherwise compromise integration. The corresponding PAGA graphs further highlighted a consistent spatial trajectory from layer 1 to layer 6, based on the domains predicted by both MaskGraphene (coordinate replacement) and MaskGraphene (coordinate transformation) (Fig.10e-f, bottom panels). In contrast, PAGA graphs derived from joint embeddings of STAligner showed misconnections across several predicted layers, failing to capture a linearly connected developmental progression. In summary, despite the challenges associated with integrating numerous adjacent consecutive ST slices, MaskGraphene successfully achieved effective integration. It generated joint embeddings that captured the geometric structure to a significant extent, revealed the spatial trajectory, and maintained strong clustering performance across individual slices.

### MaskGraphene stitches mouse brain anterior and posterior sections

Thus far, we have examined MaskGraphene’s capability to integrate adjacent consecutive tissue slices. In this section, we extended our investigation to evaluate its performance in integrating horizontally consecutive slices, focusing on two 10x Visium mouse brain sagittal sections divided into anterior and posterior portions.^10, 37^ To evaluate the integration effect, we employed the Allen Brain Atlas as reference (Fig.11a), enabling visual comparisons of MaskGraphene’s clustering results against the other six methods (Fig.11c-i). PRECAST, SPIRAL, DeepST, and GraphST were not able to detect and connect common spatial domains along the shared boundary of the anterior and posterior sections. In contrast, MaskGraphene, STAligner, and BASS successfully identified and linked common spatial domains along the shared boundary. Notably, MaskGraphene and STAligner excelled in the cerebral cortex (CTX) region, where they outperformed the other tools by accurately identifying and aligning six distinct layers across the anterior and posterior sections. Moreover, for unshared regions, both MaskGraphene and STAligner excelled in separating the caudal putamen (CP) and nucleus accumbens (ACB) and distinguishing the layers within the cerebellar cortex (CBX). Additionally, both methods effectively identified a coherent arc across two sections for CA1, CA2, and CA3. Using ground truth annotations from the anterior region (Fig.11b), we further quantified the integration performance by calculating the ARI score based on joint embeddings. MaskGraphene achieved the highest ARI score of 0.436, closely followed by STAligner (0.424). These results highlight the performance of MaskGraphene and STAligner in capturing spatial structures within complex datasets and their proficiency in integrating non-consecutive slices with batch correction.

**Fig 11.**
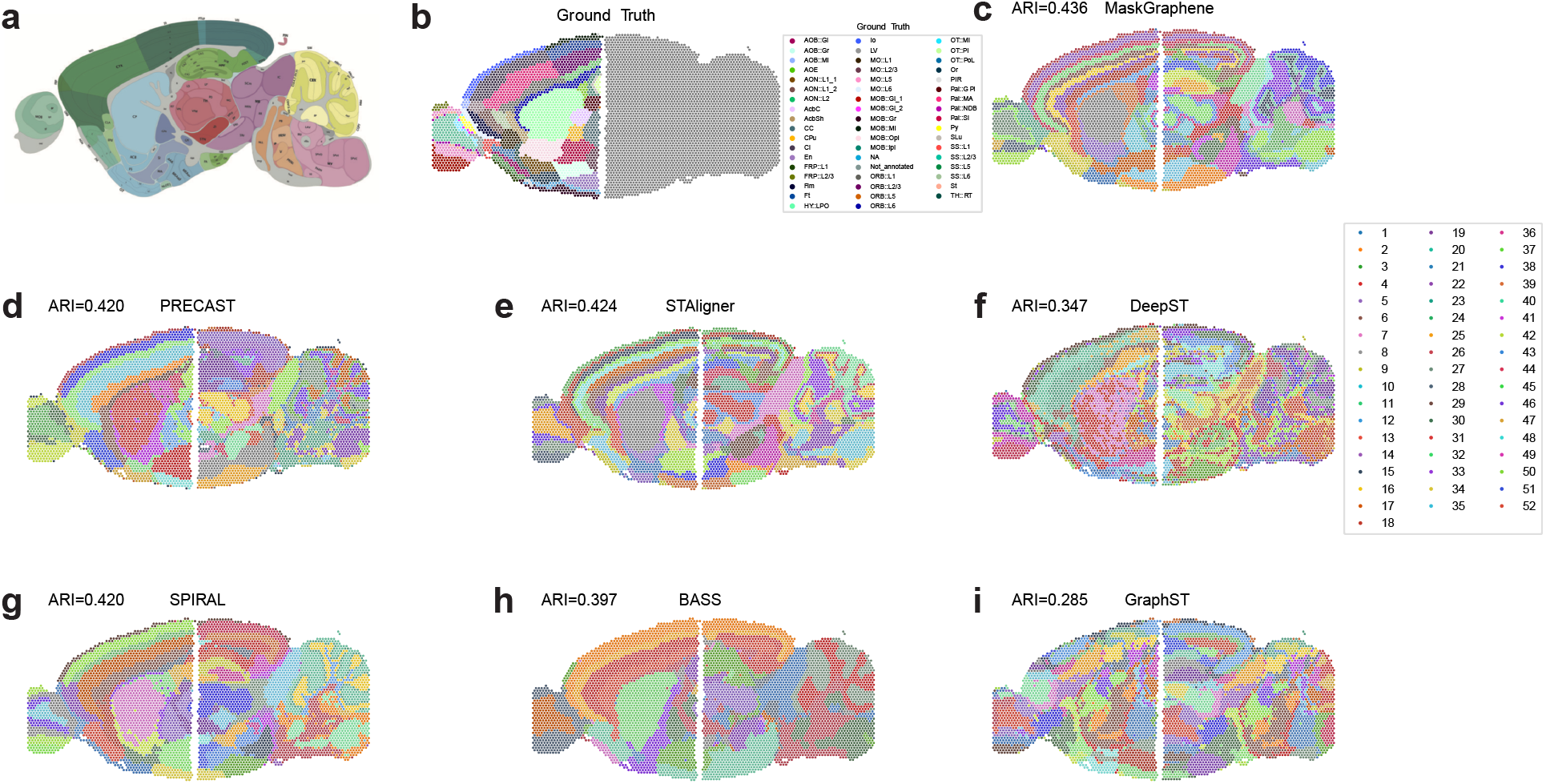
Spatial visualization plots for integration of the dataset of mouse brain sagittal sections. **(a)** The Allen brain atlas annotation for the mouse brain sagittal section. **(b)** Ground truth annotation of different regions for the anterior slice. **(c-i)** Domain identification using seven different methods on the mouse brain sagittal dataset, with ARI score evaluated for the anterior slice.

### MaskGraphene aligns tissues and organs across different developmental stages

Finally, we tested MaskGraphene’s ability to integrate two slices from different development stages for joint analysis, to study the spatiotemporal development in tissue structures during mouse organogenesis. Using Stereo-seq data from two mouse embryo slices acquired at different time points (E11.5 and E12.5),^38^ we performed an integration analysis, benchmarking MaskGraphene against STAligner. Other tools were excluded from the comparison due to memory limitations with this large dataset. As shown in Fig.12a, despite the differences in the sizes of the two slices and the presence of noticeable batch effects, both MaskGraphene and STAligner effectively harmonized the data by integrating them into a unified embedding space. They accurately detected both shared (labeled consistently across slices by the two tools and the ground truth) and developing structures unique to different time points. To evaluate clustering performance of joint embeddings, we calculated ARI scores by comparing the detected clusters against annotated ground truth labels. MaskGraphene outperformed STAligner, achieving higher ARI scores of 0.422 and 0.462 for the two slices compared to STAligner’s scores of 0.332 and 0.316.

**Fig 12.**
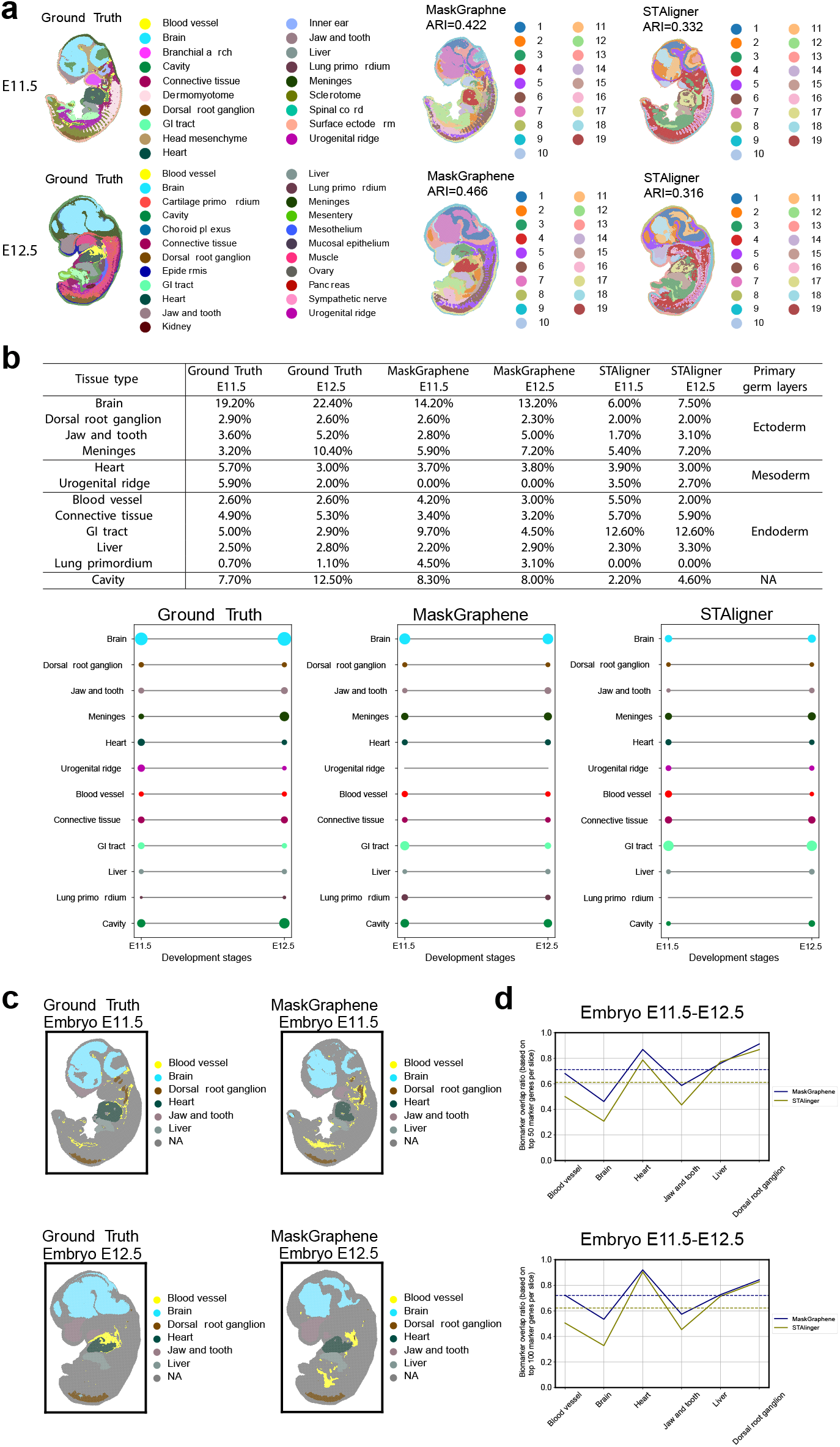
Developmental progression evaluation of spatial domain integration across embryonic developmental stages. **(a)** Spatial visualization of mouse embryos at stages E11.5 and E12.5, with ARI indicated. **(b)** Top panel: Proportion of spots for each structure relative to the total spots at time points E11.5 and E12.5, based on the ground truth and the integration results from MaskGraphene and STAligner. Bottom panel: Circles representing these proportions across time points E11.5 and E12.5. **(c)** Spatial visualizations of joint domain identification for two embryo slices based on ground truth and MaskGraphene predictions after integration. Matched colored annotations indicate several shared key tissue structures across two conditions. **(d)** Biomarker overlap ratio curves across shared tissue structures for MaskGraphene and STAligner, comparing the top *N* biomarkers identified for each integrated tissue structure after integration with the *N* ground truth marker genes. Ground truth markers are defined as the union of the top 50 (left panel) or top 100 (right panel) tissue marker genes across two slices.

To further assess the temporal development of each tissue structure during mouse organogenesis, we quantitatively analyzed the sizes of shared tissue structures across two time points by calculating the proportion of spots corresponding to each tissue structure relative to the total number of spots at each time point. For benchmarking purposes, we also computed these proportions using ground truth annotations. MaskGraphene demonstrated closer alignment with the ground truth proportions across both time points compared to STAligner. To illustrate these results, we grouped all shared tissue structures into three primary germ layers and displayed the proportions for each tool alongside the ground truth (Fig.12b, top panel). Additionally, we calculated the proportional differences across these tissue structures between each tool and the ground truth. MaskGraphene achieved a smaller difference of 45.7% and 36.3%, compared to STAligner’s 62.8% and 57.6% for E11.5 and E12.5, respectively (Supplementary Table 2). Notably, the brain, which constitutes the largest proportion of the embryo at both time points, was largely undetected by STAligner, which also missed the lung primordium. In contrast, MaskGraphene successfully detected most of the brain but failed to identify the urogenital ridge. Furthermore, MaskGraphene captured the actual proportions and temporal changes from E11.5 to E12.5 more accurately for a broader range of tissue types, including the dorsal root ganglion, jaw and tooth, and liver (Fig.12b, bottom panel). These results facilitated the reconstruction of the developmental progression of each tissue structure throughout organogenesis.

We explored whether integration analysis improves biomarker identification for these shared tissue structures. Six key structures were selected: blood vessel, brain, dorsal root ganglion, heart, jaw and tooth, and liver. To assess this, we compared the top *N* marker genes from individual slices, based on ground truth labels, with biomarkers identified from the integrated tissue structures across two embryo slices. Details of our approach are provided in the Methods section. MaskGraphene consistently achieved higher overlap ratios across nearly all tissue structures when compared to the ground truth marker genes, defined by selecting the top 50/100 tissue marker genes (Fig.12c). Notably, its performance exceeded that of STAligner, especially in the brain, where MaskGraphene exhibited significantly greater overlap ratios. These results aligned with the temporal development analysis.

In summary, the results confirm that MaskGraphene is superior in preserving spatial domains, maintaining stage-to-stage consistency, accurately representing tissue proportions across embryonic stages to some extent, and enhancing biomarker identification. These strengths are particularly valuable for developmental studies, where accurate tissue identification and cross-stage consistency are crucial.

## Discussion and Conclusion

In this work, we introduce MaskGraphene, a graph neural network that employs both self-supervised and self-contrastive training strategies to align and integrate ST data using gene expression and spatial location. To enhance interslice connectivity, MaskGraphene first performs cluster-wise alignment between consecutive slices to align spots across slices, establishing direct “hard-links” that are then used to construct the k-NN graph for the model. MaskGraphene further strengthens these connections by leveraging a graph attention autoencoder to refine low-dimensional latent embeddings through a masked self-supervised loss and triplet loss optimization. The triplet loss iteratively minimizes triplets in the latent space, acting as indirect “soft-links” that reinforce the interconnections between slices. MaskGraphene generates batch-corrected joint embeddings while preserving geometric information, thereby enhancing downstream analyses. We evaluated MaskGraphene against several state-of-the-art integration methods using a variety of real and simulated datasets, as well as multiple downstream analyses, employing diverse evaluation metrics to comprehensively assess the quality of the joint embeddings. MaskGraphene exhibited superior performance compared to all other methods.

To the best of our knowledge, existing clustering and integration tools face challenges in generating latent embeddings that effectively preserve the geometric information of original tissue slices. MaskGraphene addresses this gap, producing interpretable joint embeddings that faithfully capture and reflect the geometric structure of tissue slices. These embeddings integrate both gene expression and spatial context information to some extent, making them suitable replacements for these two modalities in various downstream analyses. For example, in our study, we utilized joint embeddings to construct a topographic map of brain slices, revealing gradients of neuronal differentiation and activity. However, the ability to generate interpretable joint embeddings is limited to certain application scenarios. For the DLPFC datasets, which exhibit layer structures across all adjacent consecutive slices, MaskGraphene’s joint embeddings effectively recover the overall slice shape, layer-wise patterns, and spatial relationships in two-dimensional UMAP space, capturing geometric information with near-perfect accuracy. In contrast, for more heterogeneous tissues, such as those in the MHypo dataset, capturing perfect geometric information becomes significantly more challenging. In the field of spatial transcriptomics, developing interpretable embeddings across all tissue slices remains a challenging yet important area of research. Such embeddings hold the potential to significantly enhance a broad spectrum of downstream analyses and provide valuable biological insights.

## Methods

### MaskGraphene workflow

The MaskGraphene workflow (Fig.1) takes as input a spot-gene expression matrix and spatial coordinates from two or more tissue slices. To strengthen interslice connections, we perform cluster-wise alignment, creating “hard-links” to unify spots within a shared coordinate system. MaskGraphene then constructs k-NN networks between spots based on their unified spatial coordinates across the slices. Leveraging a graph attention auto-encoder backbone,^23, 24^ MaskGraphene optimizes low-dimensional latent embeddings by minimizing a masked self-supervised loss^25–27^ and a triplet loss. The triplet loss is minimized by distinguishing triplets built from “soft-links” recursively through optimization. Further details are provided in the corresponding sections below. With the final joint embeddings, MaskGraphene seamlessly integrates ST slices, enabling the identification of similar spatial domains or cell types across different tissue slices. We demonstrate MaskGraphene’s effectiveness through various quantitative and qualitative downstream analyses.

### Cluster-wise alignment to generate “hard-links”

#### Spot-to-spot mapping

To enhance interslice connections, we developed an optimal transport (OT)based cluster-wise alignment method to establish reliable spot-to-spot (node-to-node) mapping (alignment) across slices, enabling the unification of all spots into a shared coordinate system for constructing the k-NN graph in MaskGraphene’s graph model. This cluster-wise process begins with an initial clustering step to identify shared clusters across slices. This is achieved by employing a graph attention autoencoder backbone, which optimizes initial joint embeddings through the minimization of a masked selfsupervised loss and a triplet loss. Details for the graph input, the model, and associated losses are provided in the next Methods sections. The resulting initial joint embeddings are then used for clustering. Once shared clusters are identified across slices, clusterwise OT-based alignment is performed. Specifically, for each pair of shared clusters (representing the same domain across each pair of slices), we solve a Gromov-Wasserstein OT problem^39^ by using the iterative conditional gradient algorithm,^40^ to generate spot-to-spot mapping.

Let (*X, D, g*) represent a slice, where *X* = [*x*_*ij*_]∈ ℕ ^*p×n*^ is a *p* genes by *n* spots transcript count matrix, *x*_*ij*_ is the transcript count for gene *i* in spot *j*, 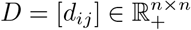 is a *n* spots by *n* spots spatial distance matrix, *d*_*ij*_ = ∥ *z*_.*i*_ − *z*_.*j*_∥ is the spatial distance between spots *i* and *j, z*_.*i*_ is the 2D coordinate of spot *i, g* = (*g*_1_, …, *g*_*n*_) is a weight vector that *g*_*i*_ *>* 0 is the weight of spot *i*.

Consider two slices (*X, D, g*) and (*X*^*′*^, *D*^*′*^, *g*^*′*^), containing *n* and *n*^*′*^ spots, respectively, with *M* shared clusters identified from the initial clustering. For each pair of shared cluster *C*^(*m*)^ between the two slices, the transport cost is minimized as follows to find a mapping between the two slices:

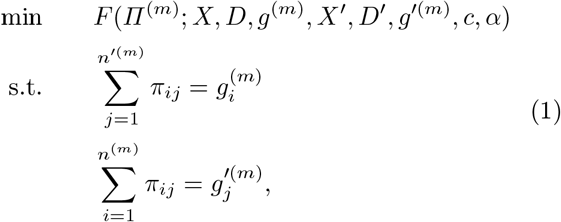

where 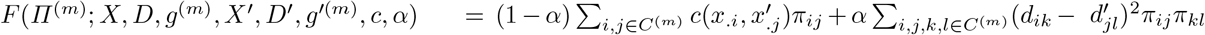, *C*^(*m*)^ contains *n*^(*m*)^ and *n*^*′*(*m*)^ spots from the two slices, respectively, *g*^(*m*)^ and *g*^*′*(*m*)^ are weight vectors of spots in *C*^(*m*)^ from the two slices, respectively. If there is no prior information about the spots, uniform distributions 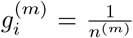 and 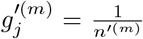 are assigned, 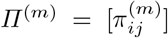 is the transport plan for cluster *C*^(*m*)^, *i* and *k* refer to spots in the first slice, *j* and *l* refer to spots in the second slice, *c* is an expression cost function, and *α* is a weight such that 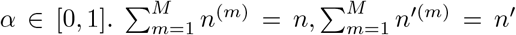 *g* = (*g*^(1)^, …, *g*^(*M*)^), *g*^*′*^ = (*g*^*′*(1)^, …, *g*^*′*(*M*)^).

The complete mapping (alignment) *?* between the two slices is obtained by combining the transport plans across all clusters:

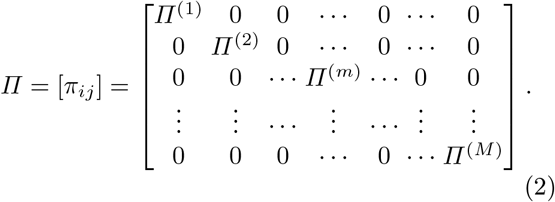

#### Coordinate system unification

After obtaining the transport plans (spot-to-spot mapping) for all clusters, referred to as “hard-links”, MaskGraphene integrates these interslice connections into its final k-NN graph model by employing three distinct strategies, each designed for specific integration scenarios.

Firstly, when the two slices (*X, D, g*) and (*X*^*′*^, *D*^*′*^, *g*^*′*^) are adjacent consecutive slices and are used for pairwise integration (e.g. pairwise DLPFC and MHypo slices along the z-axis, or mouse embryo slices from closely related developmental stages in this study), we project the second slice to the same coordinate system as the first slice after we find the mappings *?*. The following generalized weighted Procrustes problem is solved to apply for coordinate transformation:^39^

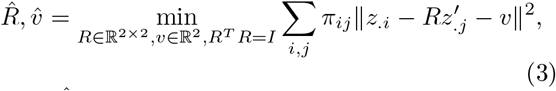

where 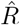 is the rotation matrix and 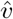 is the translation vector.

The updated spatial coordinates of spot *j* in slice (*X*^*′*^, *D*^*′*^, *g*^*′*^) are then given by:

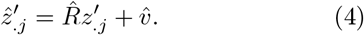

Secondly, when multiple (*>*2) consecutive ST slices (e.g., DLPF four-slice, MHypo five-slice, and MB ten-slice integration, in this study) are available for integration, achieving a robust unified coordinate system across all slices becomes increasingly challenging, particularly as the number of slices grows and geometric similarity between slices diminishes, with some adjacent slices exhibiting only partial overlap. MaskGraphene addresses this challenge by employing two strategies, coordinate replacement and coordinate transformation, to unify the coordinate system across slices. Using “hard-links” for spatial coordinate replacement, spot-to-spot mappings (alignments) are first generated for each pair of adjacent consecutive slices. The highest-confidence mappings are then iteratively applied in a cascading manner to replace coordinates, ensuring that all slices are aligned to the spatial coordinate system of the first slice. This streamlined approach enables the efficient construction of a k-NN graph that incorporates both intraslice and interslice connections across multiple slices, without requiring additional coordinate transformations.

To utilize “hard-links” to perform spatial coordinate transformation across multiple slices, we predict their coordinate systems using a mutual nearest neighbor (MNN) graph constructed from the unified spatial coordinates of the spots. The process is framed as an optimization problem guided by an objective function. To address partially overlapping slices during multi-slice integration, a penalty term is incorporated to account for the proportion of overlapping spots. Using the differential evolution optimization algorithm,^41^ we determine the optimal transformation parameters - including translation vectors and rotation angles - for each slice. These parameters are then used to transform the coordinates of each slice.

The transformation process of slice *j* is expressed mathematically as:

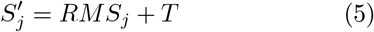

where *S*_*j*_ represents the original slice’s coordinate matrix, 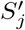is the transformed coordinate matrix, *R* is the rotation matrix:

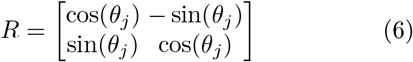

*M* is the mirroring matrix:

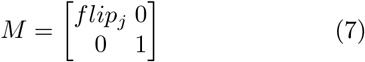

and *T* is the translation matrix:

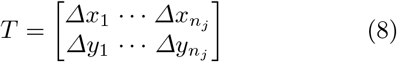

Here, *n*_*j*_ denotes the number of spots in slice *j, θ*_*j*_ is the rotation angle for slice *j*, and *flip*_*j*_ indicates whether mirroring is applied (*flip*_*j*_ = −1 for mirroring, *flip*_*j*_ = 1 otherwise).

Once the domain (cluster) information for all spots in the two adjacent slices *j*− 1 and *j* is obtained from the initial clustering, identify a rotation matrix *R* and a translation matrix *T* for slice *j* that maximize the following objective function:^30^

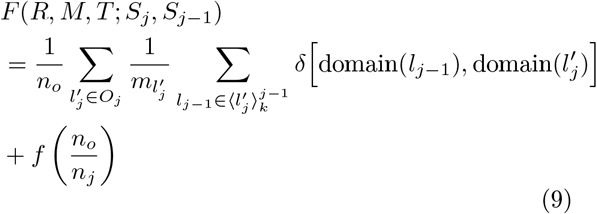

where 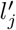 represents the overlapping spots in the transformed slice *j* whose Euclidean distance to their first mutual nearest neighbor spots in slice *j* −1 is less than a defined maximum distance. The set of those overlapping spots is denoted as *O*_*j*_, and the total number of spots in *O*_*j*_ is *n*_*o*_. By default, the maximum distance is set to the median Euclidean distance to the 2kth nearest neighbor of spots in slice *j −*1. The term 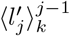 represents the set of spots in slice *j−* 1 that are *k*-nearest neighbors of spots 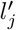, with Euclidean distance to 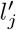 less than the maximum distance. The number of spots in 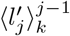 is *m*_*l*_*′*. The Kronecker delta function, *δ*(*x, y*), is used to determine whether each overlapping spot and its *k*-nearest neighboring spots belong to the same domain (cluster). *f* (*x*) = − (1− *x*)^*p*^ serves as a penalty term, where *p* controls the penalty strength. Higher values of *p* increase sensitivity to partial overlaps between adjacent slices.

We employ the differential evolution algorithm to optimize the transformation matrices {*R, M, T*}, maximizing the objective function *F* (*R, M, T* ; *S*_*j*_, *S*_*j−*1_) for adjacent slices, thereby achieving coordinate transformation across slices.

In summary, for multi-slice integration, users can select either coordinate replacement or coordinate transformation to unify the coordinate system across slices. In our study, both methods exhibited comparable performance overall; however, coordinate transformation demonstrated superior results for MB ten-slice integration.

Finally, when horizontally consecutive slices are integrated (e.g. mouse brain sagittal sections divided into anterior and posterior portions in this study), “hard-links” are directly incorporated as edges in the k-NN graph to strengthen interslice connections. Coordinate transformation or replacement is unnecessary, as these slices are horizontally connected. Subsequently, the final k-NN graph, incorporating both intraslice and interslice connections, is constructed.

It is worth noting that MaskGraphenes allows users to also utilize other state-of-the-art alignment tools to generate transport plans (spot-to-spot mappings) and unify coordinates across slices.

#### Graph input processing

MaskGraphene processes gene expression data and spatial coordinates from two or more ST slices. First, it normalizes the total count of raw gene expressions and log-transforms them using the Scanpy package.^29^ It then identifies the highly variable genes (HVGs) from all slices, and focuses on their HVGs intersection to maintain the consistency of expression features.

#### Graph model backbone

In this section, we present the mathematical formulation and details of our backbone, the graph attention autoencoder. This model comprises five main components: the encoder, graph attention mechanism, decoder, projector, and generator.

#### Encoder

The encoder generates node (spot) embeddings by aggregating information from all neighbors. We denote 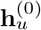 as the feature of spot *u*. The *l*^th^ encoder layer generates the embedding of spot *u* in layer *l* as follows:

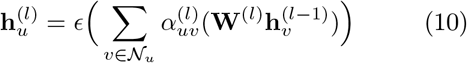

where 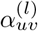 is the attention coefficient, discussed further below, *ϵ* is the activation function, *𝒩*_*u*_ denotes all the neighbors of node *u* including *u* itself, 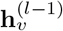 denotes the embedding of node *v* in the *l* − 1 layer, and **W**^(*l*)^ is the matrix of trainable parameters in *l*^th^ layer.

#### Graph attention mechanism

The attention coefficient 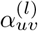 is calculated by (11) and (12) where ⊕ is the concatenation operation and *σ* is a sigmoid activation function.

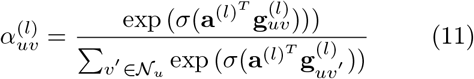

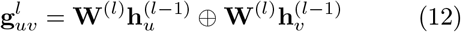

where **a**^(*l*)^ is another learnable parameter. The attention mechanism here is exploited to strengthen the connection between nodes that are represented by similar expression profiles. The attention coefficient 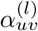 indicates the different contributions of each neighbor used in the aggregation process. The weights of edges are automatically calculated, based on node embedding.

#### Decoder

The decoder attempts to reconstruct the normalized expression profile for each spot *u* given the latent embeddings of the encoder. The *l*^th^ layer of the decoder (from the perspective of spot *u*) defined as:

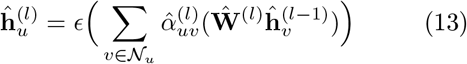

where 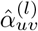 is the decoder attention coefficient which is calculated similarly as 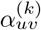 in the encoder, and Ŵ ^(*l*)^ is the matrix of trainable parameters in *l*^th^ layer.

The reconstruction loss function is introduced in the next Methods section in the context of masked self-supervised loss.

#### Projector

The projector is a multi-layer perceptron (MLP) serving as the decoder for input feature reconstruction. It consists of two layers that map the latent space to the representation space for prediction.

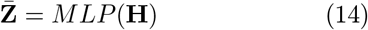

where **H** denotes the latent embedding of encoder and 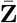 denotes the output of projector.

#### Generator

The generator has an identical structure to the aforementioned encoder and projector but with a different set of trainable parameters *ξ*_*enc*_ and *ξ*_*proj*_. It works on the unmasked graph input to produce the latent target representation 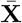, which is leveraged to train the encoder and projector network to match the output of the generator on masked nodes.

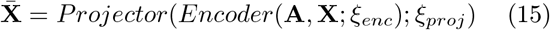

where **A** and **X** denote the adjacency matrix and input node features of the unmasked graph.

#### Masked self-supervised loss

For our graph model backbone, we introduced a selfsupervised loss to reconstruct the node features that are randomly masked from the input gene expression matrix.

Specifically, MaskGraphene perturbs the input graph by masking node features and then attempts to reconstruct the original input. A subset of input spots (e.g. 30%) is randomly selected, and their expression profiles are masked by setting them to zero. This strategy aims to enforce the reconstruction in the embedding space and regularize the node feature reconstruction.

For each node *i*, let **z**_*i*_ represent its reconstructed gene expression via the decoder. The reconstruction loss function is formulated as:

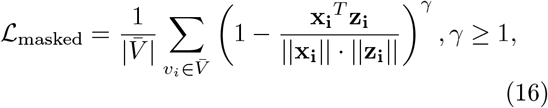

where **x**_*i*_ denotes the original normalized expression profile for spot *i*, 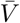 is the subset of spots with masked features in the graph, and *γ* is a scaling factor for adjusting loss. Through this scaled cosine error loss to measure the reconstruction error, the model strives to learn node embeddings under a masked autoencoder.

To further leverage the feature information, a supporting network is utilized as the target generator to produce latent prediction targets from the unmasked graph. This generator, as previously described, shares the same structure as the aforementioned encoder and projector but operates with a different set of trainable parameters. MaskGraphene simultaneously optimizes the parameters of the encoder, projector, and generator by minimizing the following loss function:

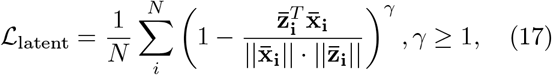

where *N* denotes all the spots in the unmasked graph, 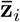 represents the output of projector from the masked graph, and 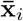 denotes the output latent targets of generator from the unmasked graph.

MaskGraphene simultaneously optimizes two objective functions with a balancing coefficient *λ*:

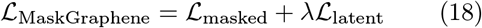

One intuition behind this is that we are enforcing the network not to smooth out those singular yet critical features from the gene expression profiles while suppressing the noises from batch effect, etc. Notably, during the initial clustering phase of the cluster-wise alignment method, this masked self-supervised loss is also utilized within the graph model to learn joint embeddings.

#### Triplet loss

To learn the final joint embeddings in the graph model, MaskGraphene leverages both “hard-links” and “soft-links” to enhance interslice connections. As mentioned previously, “hard-links” are generated using a cluster-wise method and incorporated into the k-NN graph through various strategies.

For “soft-links”, MaskGraphene employs mutual nearest neighbors (MNNs)^42^ from each pair of consecutive slices to dynamically construct spot triplets based on their embeddings during training. The interslice connections, denoted as “soft-links”, differ from “hard-links” in that they are not directly integrated into the k-NN graph. Instead, they are reinforced through the triplet loss, a method often used in singlecell RNA-seq batch correction.^21, 43^ These “soft-links” are designed to emphasize differences between dissimilar spots across slices while clustering similar spots closer together. The objective is to bring similar spots nearer and separate contrasting ones.^43^

In detail, triplet loss is based on the idea of triplets, which consist of an anchor spot **a**, a positive spot **p**, and a negative spot **n**. For each triplet (**a, p, n**), the loss is defined as follows:

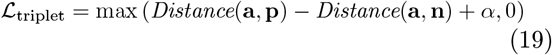

where *Distance*(**a, p**) is the Euclidean distance between the anchor spot **a** and the positive spot **p**, *Distance*(**a, n**) is the distance between the anchor spot **a** and the negative spot **n**, *α* is a margin that controls the minimum difference required between the distances of similar and dissimilar pairs. To construct triplets, we first determine mutual nearest neighbors (MNNs) across two slices. MNNs are constructed by taking the pairs of spots from distinct slices that are mutually k-nearest-neighbors.^44^ After we identify MNNs matches between two slices, we define each MNN match as the (**a, p**) pair. We further take each spot from MNN matches of one slice as an anchor spot and randomly select any spot from the other slice as a negative spot to form the (**a, n**) pair. Triplet loss encourages the model to pull the anchor and positive spots (MNNs across slices) closer in the embedding space while pushing the anchor and negative spots (dissimilar spots across slices) farther apart.

Notably, during the initial clustering phase of the cluster-wise method, “soft-links” via triplet loss are also utilized within the graph model to learn initial joint embeddings.

#### Random seed analysis

The graph model is non-deterministic and may exhibit variance, especially in clustering outcomes. To generate Adjusted Rand Index (ARI) values for clustering performance, we fixed the random seeds in MaskGraphene for all datasets. For tools that do not fix seeds, such as STAligner and DeepST, we selected the best ARI value from 10 runs for each experiment. Additionally, we evaluated the robustness of all tools by varying the random seed and performing 20 iterations for each integration experiment, with ARI box plots illustrating the results. As shown in Supplementary Fig.17, MaskGraphene consistently outperformed other tools across nearly all integration scenarios, highlighting its robustness.

#### Benchmark Datasets

We employed five ST datasets with a total of 31 slices for benchmarking, which had corresponding manual annotations shown in Supplementary Table 1.

Specifically, (1) the human DLPFC (dorsolateral prefrontal cortex) dataset, generated with 10x Visium, includes 12 human DLPFC sections with manual annotation, indicating cortical layers 1 to 6 and white matter (WM), taken from three individual samples.^28^ Each sample contains four consecutive slices (for example, slice A, B, C, and D in order). In each sample, the initial pair of slices, AB, and the final pair, CD, are directly adjacent (10 *µ*m apart), whereas the intermediate pair, BC, is situated 300 *µ*m apart. The number of spots ranges from 3,431 to 4,788, with a total of 33,538 expressed genes across all slices.

(2) The MHypo (mouse hypothalamus) dataset by MERFISH contains five manually annotated consecutive slices^17^ labeled Bregma -0.04mm (5488 cells), Bregma -0.09mm (5557 cells), Bregma -0.14mm (5926 cells), Bregma -0.19mm (5803 cells), and Bregma - 0.24mm (5543 cells). Expression measurements were taken for a common set of 155 genes. Each tissue slice includes a detailed cell annotation, identifying eight structures: third ventricle (V3), bed nuclei of the stria terminalis (BST), columns of the fornix (fx), medial preoptic area (MPA), medial preoptic nucleus (MPN), periventricular hypothalamic nucleus (PV), paraventricular hypothalamic nucleus (PVH), and paraventricular nucleus of the thalamus (PVT).

(3) The MB (mouse brain) dataset^30, 36^ by MER-FISH has a total of 33 consecutive mouse primary motor cortex tissue slices with similar shapes, which can be used for 3D reconstruction. We selected the first 10 consecutive slices for benchmarking in this study. Region annotations include the six layers (L1-L6) and white matter (WM). The number of spots ranges from 2,033 to 5,624, with a total of 254 expressed genes.

(4) The dataset of mouse brain sagittal sections, generated with 10x Visium, includes two slices of the anterior and posterior mouse brain. The number of spots is 2695 and 3353 for two slices, respectively, with both containing 32,285 expressed genes. Only the anterior section includes annotations.^10, 37^ and (5) The mouse embryo dataset by Stereo-seq has over 50 slices, and the slices at two different time points E11.5 and E12.5 were used in our experiments. The number of spots is 30,124 and 51,365 for slices E11.5 and E12.5, respectively, with 26,854 and 27,810 captured and expressed genes. These data are from a large stereo-seq project called MOSTA:^38^ Mouse Organogenesis Spatiotemporal Transcriptomic Atlas by BGI.

#### Simulated Datasets

Due to the limited availability of benchmark datasets for evaluating spot-to-spot alignment accuracy, we employed our recent ST simulation method^11^ to generate five simulated 10x Visium datasets for this purpose. Specifically, we used the DLPFC 151673 slice as the reference and created new slices by altering the spatial coordinates of the reference through rotation. To further simulate real-world scenarios, we adjusted the number of spots in each slice by removing spots that no longer aligned with the grid coordinates after rotation. To preserve the fidelity of real 10x Visium data, we arranged the tissue spots in a hexagonal grid pattern rather than a rectangular one. By applying rotations with different degrees, we generated slices with varying overlap ratios (20%, 40%, 60%, 80%, and 100%) relative to the reference slice.

### Quantitative metrics

#### Adjusted Rand Index (ARI)

ARI measures the similarity between two data clusterings by assessing the concordance of pairwise data point groupings. It evaluates whether pairs of points are assigned to the same cluster or different clusters in the two clusterings. The ARI ranges from -1 to 1, where 1 represents perfect agreement, 0 indicates clustering that is no better than random, and negative values suggest disagreement between the clusterings.

#### Layer-wise alignment accuracy

This metric^11^ is based on the hypothesis that aligned spots from adjacent consecutive slices are likely to belong to the same spatial domain or cell type. Joint spot embeddings from each method are used to align (anchor) spots from the first slice to corresponding (aligned) spots on the second slice for each slice pair. Spot proximity for alignment is determined using Euclidean distance. Alignment accuracy is calculated as the proportion of anchor spots in the first slice that are aligned to spots in the second slice belonging to the same spatial domain or cell type. For the DLPFC slices, which exhibit a unique layered structure, this metric is further designed to assess whether anchor and aligned spots belong to the same layer (layer shift = 0) or they belong to different layers (layer shift = 1 to 6). This design ensures the metric accurately captures alignment performance within the unique context of layered structures.

#### Spot-to-spot matching ratio

This metric^11^ quantifies the accuracy of alignment between corresponding spots on adjacent tissue slices. It is defined as the ratio of anchor spots in the first slice to aligned spots in the second. An optimal integration method for two adjacent consecutive slices would ideally result in a 1:1 ratio, indicating perfect alignment fidelity.

#### Spot-to-spot alignment accuracy

To assess spot alignment across slices, this metric^11^ is applied to simulated datasets with known ground truth, measuring spot-wise alignment accuracy. It is defined as the percentage of anchor spots in the simulated slice that are correctly matched to their corresponding aligned spots in the reference slice.

#### Pearson correlation coefficient

To assess the alignment of spots, we computed the Pearson correlation coefficient for matched spots identified using known ground truth in simulated datasets. This coefficient measures the strength and direction of the linear relationship between the embeddings for the matched spots, with values closer to 1 indicating a stronger linear alignment.

#### Integration local inverse Simpson’s index (iLISI)

The iLISI metric^15^ is used to assess the quality of batch mixing following data integration in spatial transcriptomics or single-cell RNA sequencing. It quantifies the extent to which the integrated data achieves a balance between preserving biological variability across cell types and promoting effective batch mixing. iLISI ranges from 0 to 1. A higher iLISI score indicates more even mixing of spots or cells from different batches, reflecting the effectiveness of the integration method in mitigating batch effects while preserving the underlying biological structure of the data.

#### Isometry correlation and Procrustes dissimilarity

In this study, geometric information preservation refers to maintaining the spatial structure and relationships between spots or cells when projecting into latent embedding space after ST integration.

This involves preserving both isometric relationships and the overall similarity in spatial arrangements across datasets. To assess the quality of preservation, we used two metrics, Isometry correlation and Procrustes dissimilarity.

Isometry correlation evaluates whether the relative distances between points in the original spatial data are preserved in the integrated or embedded space. Ideally, points that are close in the original space should remain close after integration, while points that are far apart should maintain proportional distances. Isometry correlation ranges from 0 to 1. A higher Isometry correlation indicates that the intrinsic spatial organization of the data is well-preserved, supporting accurate biological interpretation. Procrustes dissimilarity, on the other hand, compares two datasets by applying transformations such as scaling, rotation, and reflection to align one dataset as closely as possible with the other. This metric quantifies how well the original tissue geometry is preserved in the embedding space. Procrustes dissimilarity ranges from 0 to 1. A smaller Procrustes dissimilarity value indicates better alignment, reflecting that the spatial relationships between spots or cells have been well-preserved across datasets.

### Qualitative analysis

#### Visualization of aligned, misaligned, and unaligned spots from pairwise integration and alignment

To visually evaluate the quality of joint spot embeddings, we aligned each spot on the first slice (the “anchor” spot) with a corresponding spot on the second slice (the “aligned” spot) based on their joint latent embeddings generated by each integration method, using Euclidean distance. If the aligned spot belonged to the same spatial domain or cell type as the anchor spot based on ground truth labels, both spots were classified as “aligned” and marked in orange (Fig. 3a-b). If the aligned spot differed in spatial domain or cell type from the anchor spot, both were classified as “mis-aligned” and marked in blue. Finally, spots on the second slice that were not aligned to any spot on the first slice were classified as “unaligned” and marked in green (Fig. 3a-b).

#### Spatial visualization of joint domain identification through clustering results after integration

For all datasets we used, we compared the identified domains across slices after integration with the ground truth labels. For the dataset of mouse brain sagittal sections, we further compared the identified domains after integration with the Allen Brain atlas through visualization. Additionally, we assessed the consistency of regions across the fissure between the anterior and posterior sections. Greater similarity to the atlas and improved regional coherence indicate superior integration performance.

#### Visualization of UMAP plot for joint embeddings

The majority of integration techniques focus on embedding spots into a low-dimensional latent space, which can often be difficult to interpret intuitively. To improve understanding of the distribution within the latent space, we applied UMAP for dimensionality reduction, projecting the spot embeddings into two dimensions. An effective UMAP visualization of latent embeddings should display patterns that reflect the characteristics of the original data while clearly delineating spatial domains or cell types.

#### Partition-based graph abstraction (PAGA) analysis

PAGA is a computational framework primarily used in single-cell genomics to infer and visualize the connectivity structure of cell clusters or partitions, often along developmental trajectories.^45^ It combines graph-based clustering with a simplified abstract representation of the relationships between cell groups, providing insights into the hierarchical or sequential organization of data. In this study, we used joint embeddings generated by the integration method as input for Scanpy to construct the PAGA graph, uncovering the spatial trajectory of cortical layers in both the DLPFC and MB datasets.

#### Biomarker identification analysis

To evaluate whether integration enhances biomarker identification, we compared domain-specific marker genes identified from individual slices based on ground truth labels with biomarkers derived from each integrated domain across four slices. Initially, predicted domains from the integration tool were aligned with ground truth labels to annotate the domains, and spots corresponding to each domain were extracted from all integrated slices. Biomarkers for each integrated domain were then identified using Scanpy. To create ground truth domain marker genes for comparison, we applied Scanpy to identify biomarkers for each domain in individual slices based on ground truth labels, selecting the top 50 or 100 genes per domain per slice. The union of these top genes, denoted as *N*, across all slices was compiled to form the final set of ground truth domain marker genes for validation. Finally, we computed the overlap ratio between the top *N* biomarkers identified for each integrated domain and the *N* ground truth domain marker genes. We performed this analysis for both DLPFC and mouse embryo datasets.

#### Isodepth gradient analysis

To validate the quality of joint embedding, we employed GASTON,^3^ an unsupervised and interpretable deep learning algorithm designed to generate a topographic map of brain tissue slices, and reveal gradients of neuronal differentiation and activity. GASTON constructs this topographic map by leveraging an innovative 1-dimensional coordinate system, termed isodepth, to define spatial organization within the tissue. Analogous to contour lines on a geographic map, isodepth contours delineate distinct spatial domains and quantify gradual expression changes across these regions. To estimate the isodepth *d*, GASTON solves a maximum likelihood problem by parameterizing the functions *d* : ℝ^2^ → ℝ and *h* = (*h*_1_, …, *h*_*G*_) : ℝ → ℝ^*G*^ using neural networks with weights *θ* and *θ*^*′*^, respectively.

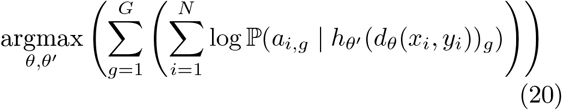

where *a*_*i,g*_ represents the UMI count of gene *g* in spot *i*. Solving Equation (15) is equivalent to training a single multi-layer perceptron with a 2D input dimension and an output dimension of G (where G denotes the total number of genes). One of the hidden layers in the network contains a single hidden neuron, whose value corresponds to the estimated isodepth. Consequently, the isodepth is represented as an interpretable hidden layer within the neural network. As defined in the orginal paper, the Spatial Topography Problem (STP) is approximated and solved by utilizing the proposed objective function in conjunction with the neural networks.

To account for both continuous variations in expression within or across spatial domains and abrupt discontinuities at the boundaries of adjacent spatial domains, the estimated isodepth from Equation (15) is employed to estimate the piecewise linear function ĥ by solving the following optimization problem.

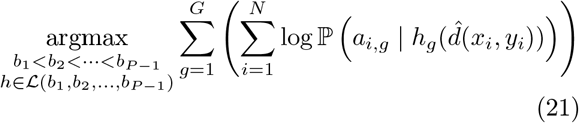

where *ℒ* () is the set of piecewise linear functions with breakpoint *b*_1_, *b*_2_, …, *b*_*P −*1_. The piecewise linear expression functions *h*_*g*_(*θ*) reveal both discontinuities in expression and continuous variation within a domain, or intradomain variation.

We compared the joint embedding of MaskGraphene with STAligner and GraphST under two different scenarios when running GASTON. By default, GASTON uses the two-dimensional X,Y coordinates of all spots and the principal components (PCs) of the expression data from a single slice to fit the above models. In the first experimental scenario, we replaced the PCs with the joint embedding from each integration method while retaining the original X,Y coordinates as input to GASTON. In the second experimental scenario, we replaced the original X,Y coordinates with the two-dimensional UMAP coordinates, which represent a low-dimensional embedding that preserves the structure of the high-dimensional data, and replaced the PCs with the joint embedding from each integration method as input to GASTON.

#### Computation platform

All experiments were conducted on a computer server equipped with Intel Xeon W-2195 CPUs, running at 2.3 GHz, featuring 25MB of L3 cache and 36 CPU cores. The server was configured with 256GB of DDR4 memory operating at 2,666MHz.

For GPU configurations, the same server was used, equipped with four Quadro RTX A6000 cards, each providing 48GB of memory and 4608 CUDA cores.

## Acknowledgements

This work was supported by the NIGMS Maximizing Investigators’ Research Award (MIRA) R35 GM146960, Young Collaborative Research Grant (C2004-23Y), and HMRF (11221026).

## Data and code availability

All code, tutorials, evaluation scripts, and related data files are freely available on GitHub^46^ https://github.com/maiziezhoulab/MaskGraphene. All data and the corresponding annotation can be downloaded from https://benchmarkst-reproducibility. readthedocs.io/en/latest/Data% 20availability.html and are described in Supplementary Table 1 with their sources. Dataset 1 consists of 12 human DLPFC sections, available at http://research.libd.org/spatialLIBD/ with manual annotation.^47^ Dataset 2^48^ includes five slices from the mouse hypothalamus available at https://datadryad.org/stash/dataset/doi: 10.5061/dryad.8t8s248 with annotation.^17^ Dataset 3^36^ contains 33 consecutive mouse cerebral cortex tissue slices with similar shapes at https://zenodo.org/records/8167488 with annotation.^30^ Dataset 4^49^ includes two slices of anterior and posterior mouse brain available at https://www.10xgenomics.com/ with annotation.^37^ Dataset 5 is the Embryo dataset sequenced by Stereo-seq from the MOSTA project at https://db.cngb.org/stomics/mosta/resource/ with annotation.^38^ The simulation data is deposited in Zenodo https://zenodo.org/records/10800745.^50^

## References

1. Perrimon, N., Pitsouli, C. and Shilo, B.-Z. (2012). Signaling mechanisms controlling cell fate and embryonic patterning. Cold Spring Harbor perspectives in biology, 4(8):a005975.

2. Zhu, J., Wang, Y., Chang, W. Y., Malewska, A., Napolitano, F., Gahan, J. C., Unni, N., Zhao, M., Yuan, R., Wu, F. et al. (2024). Mapping cellular interactions from spatially resolved transcriptomics data. Nature Methods, 1–13.

3. Chitra, U., Arnold, B. J., Sarkar, H., Ma, C., Lopez-Darwin, S., Sanno, K. and Raphael, B. J. (2024). Mapping the topography of spatial gene expression with interpretable deep learning. In International Conference on Research in Computational Molecular Biology 368– 371, Springer.

4. Marx, V. (2021). Method of the Year: spatially resolved transcriptomics. Nature Methods, 18(1):9–14.

5. Dong, K. and Zhang, S. (2022). Deciphering spatial domains from spatially resolved transcriptomics with an adaptive graph attention auto-encoder. Nature communications, 13(1):1–12.

6. Chen, J., McSwiggen, D. and Ünal, E. (2018). Single molecule fluorescence in situ hybridization (smFISH) analysis in budding yeast vegetative growth and meiosis. JoVE (Journal of Visualized Experiments), None (135):e57774.

7. Wang, X., Allen, W. E., Wright, M. A., Sylwestrak, E. L., Samusik, N., Vesuna, S., Evans, K., Liu, C., Ramakrishnan, C., Liu, J. et al. (2018). Three-dimensional intact-tissue sequencing of single-cell transcriptional states. Science, 361(6400):eaat5691.

8. Moffitt, J. R. et al. (2016). High-throughput single-cell gene-expression profiling with multiplexed error-robust fluorescence in situ hybridization. Proceedings of the National Academy of Sciences, 113(39):11046–11051.

9. Rodriques, S. G. et al. (2019). Slide-seq: A scalable technology for measuring genome-wide expression at high spatial resolution. Science, 363(6434):1463–1467.

10. Ståhl, P. L., Salmén, F., Vickovic, S., Lundmark, A., Navarro, J. F., Magnusson, J., Giacomello, S., Asp, M., Westholm, J. O., Huss, M. et al. (2016). Visualization and analysis of gene expression in tissue sections by spatial transcriptomics. Science, 353(6294):78–82.

11. Hu, Y., Xie, M., Li, Y., Rao, M., Shen, W., Luo, C., Qin, H., Baek, J. and Zhou, X. M. (2024). Benchmarking clustering, alignment, and integration methods for spatial transcriptomics. Genome Biology, 25(1):212.

12. Zhou, X. (2022). Graphing cell relations in spatial transcriptomics. Nature Computational Science, 2(6):354–355.

13. Hu, Y., Zhao, Y., Schunk, C. T., Ma, Y., Derr, T. and Zhou, X. M. (2023). ADEPT: Autoencoder with differentially expressed genes and imputation for robust spatial transcriptomics clustering. Iscience, 26(6).

14. Xu, C., Jin, X., Wei, S., Wang, P., Luo, M., Xu, Z., Yang, W., Cai, Y., Xiao, L., Lin, X. et al. (2022). DeepST: identifying spatial domains in spatial transcriptomics by deep learning. Nucleic Acids Research, 50(22):e131–e131.

15. Korsunsky, I., Millard, N., Fan, J., Slowikowski, K., Zhang, F., Wei, K., Baglaenko, Y., Brenner, M., Loh, P.-r. and Raychaudhuri, S. (2019). Fast, sensitive and accurate integration of single-cell data with Harmony. Nature methods, 16(12):1289–1296.

16. Stuart, T., Butler, A., Hoffman, P., Hafemeister, C., Papalexi, E., Mauck, W. M., Hao, Y., Stoeckius, M., Smibert, P. and Satija, R. (2019). Comprehensive integration of single-cell data. Cell, 177(7):1888–1902.

17. Li, Z. and Zhou, X. (2022). BASS: multi-scale and multi-sample analysis enables accurate cell type clustering and spatial domain detection in spatial transcriptomic studies. Genome biology, 23(1):168.

18. Liu, W., Liao, X., Luo, Z., Yang, Y., Lau, M. C., Jiao, Y., Shi, X., Zhai, W., Ji, H., Yeong, J. et al. (2023). Probabilistic embedding, clustering, and alignment for integrating spatial transcriptomics data with PRECAST. Nature communications, 14(1):296.

19. Long, Y., Ang, K. S., Li, M., Chong, K. L. K., Sethi, R., Zhong, C., Xu, H., Ong, Z., Sachaphibulkij, K., Chen, A. et al. (2023). Spatially informed clustering, integration, and deconvolution of spatial transcriptomics with GraphST. Nature Communications, 14(1):1155.

20. Guo, T., Yuan, Z., Pan, Y., Wang, J., Chen, F., Zhang, M. Q. and Li, X. (2023). SPIRAL: integrating and aligning spatially resolved transcriptomics data across different experiments, conditions, and technologies. Genome Biology, 24(1):241.

21. Zhou, X., Dong, K. and Zhang, S. (2023). Integrating spatial transcriptomics data across different conditions, technologies and developmental stages. Nature Computational Science, 3(10):894–906.

22. Duan, B., Chen, S., Cheng, X. and Liu, Q. (2024). Multi-slice spatial transcriptome domain analysis with SpaDo. Genome Biology, 25(1):73.

23. Veličković, P., Cucurull, G., Casanova, A., Romero,, Lio, P. and Bengio, Y. (2017). Graph attention networks. arXiv preprint 1710.10903,.

24. Salehi, A. and Davulcu, H. (2019). Graph attention auto-encoders. arXiv preprint 1905.10715,.

25. Hou, Z., Liu, X., Cen, Y., Dong, Y., Yang, H., Wang, C. and Tang, J. (2022). Graphmae: Self-supervised masked graph autoencoders. In Proceedings of the 28th ACM SIGKDD Conference on Knowledge Discovery and Data Mining 594–604, None.

26. Hou, Z., He, Y., Cen, Y., Liu, X., Dong, Y., Kharlamov, E. and Tang, J. (2023). GraphMAE2: A Decoding-Enhanced Masked Self-Supervised Graph Learner. In Proceedings of the ACM Web Conference 2023 737–746, None.

27. He, K., Chen, X., Xie, S., Li, Y., Dollár, P. and Girshick, R. (2022). Masked autoencoders are scalable vision learners. In Proceedings of the IEEE/CVF conference on computer vision and pattern recognition 16000– 16009, None.

28. Maynard, K. R., Collado-Torres, L., Weber, L. M., Uytingco, C., Barry, B. K., Williams, S. R., Catallini, J. L., Tran, M. N., Besich, Z., Tippani, M. et al. (2021). Transcriptome-scale spatial gene expression in the human dorsolateral prefrontal cortex. Nature neuroscience, 24(3):425–436.

29. Wolf, F. A., Angerer, P. and Theis, F. J. (2018). SCANPY: large-scale single-cell gene expression data analysis. Genome biology, 19(1):1–5.

30. Xu, H., Lin, J., Wang, S., Fang, M., Luo, S., Chen, C., Wan, S., Wang, R., Tang, M., Xue, T. et al. (2023). SPACEL: characterizing spatial transcriptome architectures by deep-learning. None,.

31. Micheva, K. D., Wolman, D., Mensh, B. D., Pax, E., Buchanan, J., Smith, S. J. and Bock, D. D. (2016). A large fraction of neocortical myelin ensheathes axons of local inhibitory neurons. elife, 5:e15784.

32. Nelson, S. B. and Valakh, V. (2015). Excitatory/inhibitory balance and circuit homeostasis in autism spectrum disorders. Neuron, 87(4):684–698.

33. Simons, M. and Nave, K.-A. (2016). Oligodendrocytes: myelination and axonal support. Cold Spring Harbor perspectives in biology, 8(1):a020479.

34. Harris, K. D. and Shepherd, G. M. (2015). The neocortical circuit: themes and variations. Nature neuroscience, 18(2):170–181.

35. Fraley, C., Raftery, A. E., Murphy, T. B. and Scrucca, L. (2012). mclust version 4 for R: normal mixture modeling for model-based clustering, classification, and density estimation. Technical report Technical report.

36. Zhang, M., Eichhorn, S. W., Zingg, B., Yao, Z., Cotter, K., Zeng, H., Dong, H. and Zhuang, X. (2021). Spatially resolved cell atlas of the mouse primary motor cortex by MERFISH.

37. Zeng, Y., Yin, R., Luo, M., Chen, J. et al. (2022). Deciphering Spatial Domains by Integrating Histopathological Image and Tran-scriptomics via Contrastive Learning. biorxiv,.

38. Chen, A., Liao, S., Cheng, M., Ma, K., Wu, L., Lai, Y., Qiu, X., Yang, J., Xu, J., Hao, S. et al. (2022). Spatiotemporal transcriptomic atlas of mouse organogenesis using DNA nanoball-patterned arrays. Cell, 185(10): 1777–1792.

39. Zeira, R., Land, M., Strzalkowski, A. and Raphael, B. J. (2022). Alignment and integration of spatial transcriptomics data. Nature Methods, 19(5):567–575.

40. Titouan, V., Courty, N., Tavenard, R. and Flamary, R. (2019). Optimal transport for structured data with application on graphs. In International Conference on Machine Learning 6275–6284, PMLR.

41. Storn, R. and Price, K. (1997). Differential evolution– a simple and efficient heuristic for global optimization over continuous spaces. Journal of global optimization, 11:341–359.

42. Haghverdi, L., Lun, A. T., Morgan, M. D. and Marioni, J. C. (2018). Batch effects in single-cell RNAsequencing data are corrected by matching mutual nearest neighbors. Nature biotechnology, 36(5):421–427.

43. Yu, X., Xu, X., Zhang, J. and Li, X. (2023). Batch alignment of single-cell transcriptomics data using deep metric learning. Nature Communications, 14(1):960.

44. Hie, B., Bryson, B. and Berger, B. (2019). Efficient integration of heterogeneous single-cell transcriptomes using Scanorama. Nature biotechnology, 37(6):685–691.

45. Wolf, F. A., Hamey, F. K., Plass, M., Solana, J., Dahlin, J. S., Göttgens, B., Rajewsky, N., Simon, L. and Theis, F. J. (2019). PAGA: graph abstraction reconciles clustering with trajectory inference through a topology preserving map of single cells. Genome biology, 20(1): 1–9.

46. Hu, Y. and Zhou, X. M. (2024). MaskGraphene: Advancing joint embedding, clustering, and batch correction for spatial transcriptomics using graph-based selfsupervised learning.

47. Pardo, B., Spangler, A., Weber, L. M., Page, S. C., Hicks, S. C., Jaffe, A. E., Martinowich, K., Maynard, K. R. and Collado-Torres, L. (2022). spatialLIBD: an R/Bioconductor package to visualize spatially-resolved transcriptomics data.

48. Moffitt, J. R., Bambah-Mukku, D., Eichhorn, S. W., Vaughn, E., Shekhar, K., Perez, J. D., Rubinstein, N. D., Hao, J., Regev, A., Dulac, C. et al. (2018). Molecular, spatial, and functional single-cell profiling of the hypothalamic preoptic region.

49. 10x Genomics (2022). Mouse Brain Serial Section 2 (Sagittal-Anterior).

50. Hu, Y. and Zhou, X. M. (2024). DLPFC 151673 simulated data.

