## Appendix for "MaskGraphene: an advanced framework for interpretable joint representation for multi-slice, multi-condition spatial transcriptomics"

Hu et al.

11 Spatial visualization of joint domain identification after four-slice integration (151507-151508-151509-151510) . . 16

12 **Spatial visualization of joint domain identification after MHypo Bregma -0.19 - -0.24 pair-wise integration** . . . . 17
13 **Spatial visualization of joint domain identification after MHypo five-slice integration** . . . . . . . . . . . . . . . . . . 18
14 **UMAP, PAGA, and biomarker analysis of MaskGraphene (coordinate transformation) after multi-slice integration** 19
15 **Spatial visualization of joint domain identification and ARI boxplots by MaskGraphene (coordinate transformation) after multi-slice integration** . . . . . . . . . . . . . 20
16 **UMAP plots of low dimensional joint embedding on the MB dataset** . . . . . . . . . . . . . . . . . . . . . . . . . . . . . 21
17 **ARI box plots of clustering results after integration with varying random seeds across different datasets and methods** . . . . . . . . . . . . . . . . . . . . . . . . . . . . . . . . . . 22

### 1 Supplementary Tables

| Algorithm | Language | Resource | Output | Method | Link |
|---|---|---|---|---|---|
| MaskGraphene | Python | Hu et al. 2024 | Clustering Labels<br>Embedding | Graph Autoencoder<br>Self-supervised Learning<br>Cluster-wise Alignment | `https://github.com/maiziezhoulab/MaskGraphene` |
| SpaDo | R | Duan et al. 2024[1] | Clustering Labels<br>Embedding | kNN<br>multi-slice clustering | `https://github.com/bm2-lab/SpaDo` |
| SPIRAL | Python | Guo et al. 2023[2] | Clustering Labels<br>Refined Coordinates<br>Embedding | GraphSAGE<br>Optimal Transport | `https://github.com/guott15/SPIRAL` |
| STAligner | Python | Zhou et al. 2023[3] | Clustering Labels<br>Embedding | Graph Autoencoder<br>Attention Mechanism<br>Triplet Loss | `https://github.com/zhanglabtools/STAligner` |
| PRECAST | R | Liu et al. 2023[4] | Clustering Labels<br>Embedding | Gaussian Mixture Model<br>Discrete Hidden<br>Markov Random Field | `https://github.com/cran/PRECAST` |
| BASS | R | Li et al. 2022[5] | Clustering Labels | Bayesian Analysis<br>Multi-sample Analysis | `https://github.com/zhengli09/BASS` |
| DeepST | Python | Xu et al. 2022[6] | Clustering Labels<br>Embedding | Data Augmentation<br>Variational Autoencoder | `https://github.com/JiangBioLab/DeepST` |
| GraphST | Python | Long et al. 2022[7] | Clustering Labels<br>Embedding | Graph Neural Network<br>Contrastive Learning | `https://github.com/JinmiaoChenLab/GraphST` |

| ST Dataset | Abbreviations | ST protocol | Spots/Genes | Num. of used slices | Source |
|---|---|---|---|---|---|
| Dataset 1: Human Dorsal Lateral Prefrontal Cortex data [8] | DLPFC | 10x Visium | 3431-4788/33538 | 12 | `http://spatial.libd.org/spatialLIBD/` |
| Dataset 2: Mouse Hypothalamus [9] | MHypo | MERFISH | 5488-5926/155 | 5 | `https://datadryad.org/stash/dataset/doi:10.5061/dryad.8t8s248` |
| Dataset 3: Mouse Brain [10] | MB | MERFISH | 2033-5624/254 | 10 | `https://zenodo.org/records/8167488` |
| Dataset 4: Mouse Brain Section 2 Sagittal Anterior and Posterior [11] | MB2SA | 10x Visium | 2695,3353/32285 | 2 | `https://www.10xgenomics.com/resources/datasets/mouse-brain-serial-section-2-sagittal-anterior-1-standard` |
| Dataset 5: MOSTA Embryo [12] | Embryo | Stereo-seq | 30124,51365/26854,27810 | 2 | `https://db.cngb.org/stomics/mosta/resource/` |

Supplementary Table 1: **Benchmark tools and real datasets.Top panel**: Summary of benchmarked methods used in this work. The tool's programming language, resource, output, the general method by each tool, and tool links are shown in the table. **Bottom panel**: Overview of the real datasets benchmarked in this study. The datasets' abbreviations in the study, ST protocol, the range of number of spots (cells) and genes, number of slices, and source link for each dataset are shown in the table.

| Tissue type | Ground Truth E11.5 | Ground Truth E12.5 | MaskGraphene E11.5 | MaskGraphene E12.5 | STAligner E11.5 | STAligner E12.5 | Primary germ layers |
|---|---|---|---|---|---|---|---|
| **Brain** | 19.20% | 22.40% | 14.20% (-26.04%) | 13.20% (-41.07%) | 6.00% (-68.75%) | 7.50% (-66.52%) | Ectoderm |
| **Dorsal root ganglion** | 2.90% | 2.60% | 2.60% (-10.34%) | 2.30% (-11.54%) | 2.00% (-31.03%) | 2.00% (-23.08%) | |
| **Jaw and tooth** | 3.60% | 5.20% | 2.80% (-22.22%) | 5.00% (-3.85%) | 1.70% (-52.78%) | 3.10% (-40.38%) | |
| **Meninges** | 3.20% | 10.40% | 5.90% (+84.37%) | 7.20% (-30.77%) | 5.40% (+68.75%) | 7.20% (-30.77%) | |
| **Heart** | 5.70% | 3.00% | 3.70% (-35.09%) | 3.80% (+26.67%) | 3.90% (-31.58%) | 3.00% (0.00%) | Mesoderm |
| **Urogenital ridge** | 5.90% | 2.00% | 0.00% (-100.00%) | 0.00% (-100.00%) | 3.50% (-40.68%) | 2.70% (+35.00%) | |
| **Blood vessel** | 2.60% | 2.60% | 4.20% (+61.54%) | 3.00% (+15.38%) | 5.50% (+111.54%) | 2.00% (-23.08%) | Endoderm |
| **Connective tissue** | 4.90% | 5.30% | 3.40% (-30.61%) | 3.20% (-39.62%) | 5.70% (+16.33%) | 5.90% (+11.32%) | |
| **GI tract** | 5.00% | 2.90% | 9.70% (+94.00%) | 4.50% (+55.17%) | 12.60% (+152.00%) | 12.60% (+334.48%) | |
| **Liver** | 2.50% | 2.80% | 2.20% (-12.00%) | 2.90% (+3.57%) | 2.30% (-8.00%) | 3.30% (+17.86%) | |
| **Lung primordium** | 0.70% | 1.10% | 4.50% (+542.86%) | 3.10% (+181.82%) | 0.00% (-100.00%) | 0.00% (-100.00%) | |
| **Cavity** | 7.70% | 12.50% | 8.30% (+7.79%) | 8.00% (-36.00%) | 2.20% (-71.43%) | 4.60% (-63.20%) | NA |

Supplementary Table 2: **Proportional distribution of tissue structures at each embryo stage.**Proportion of spots for each structure relative to the total spots at time points E11.5 and E12.5, based on the ground truth and the integration results from MaskGraphene and STAligner. The ratios in parentheses represent the proportional differences of each tool compared to the ground truth.

#### 2 Supplementary Figures

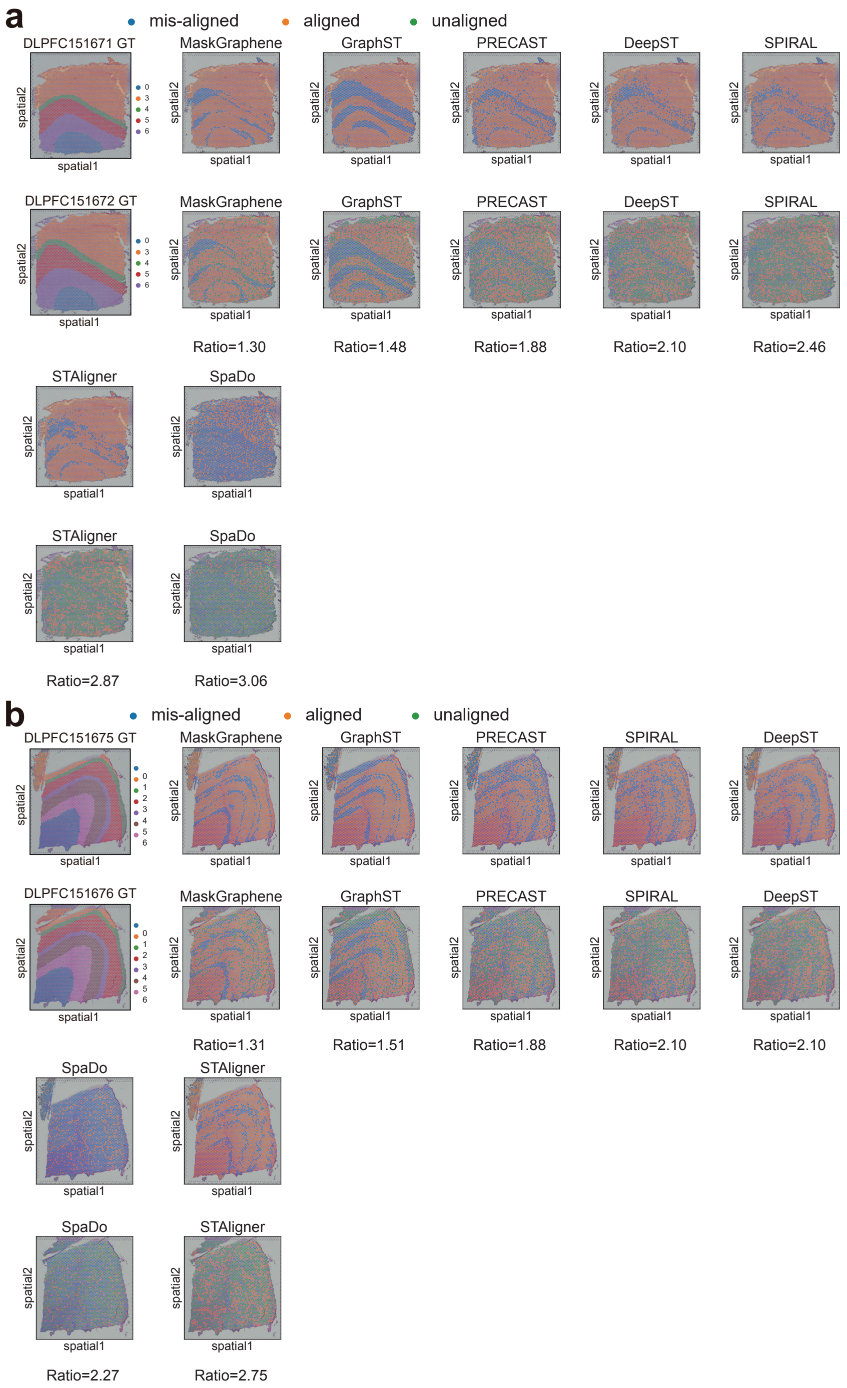

Supplementary Figure 1: **Visualization plots for alignment-misalignment-unalignment and spot-to-spot mapping ratio on DLPFC 151671-151672 pair. (a-b)** Visualization plots displaying aligned spots, misaligned spots, and unaligned spots during the alignment process. The anchor spot from the first (top) slice is aligned to the corresponding spots on the second (bottom) slice for DLPFC 151671-151672 pair **(a)** and DLPFC 151675-151676 pair **(b)**. The first slice pair illustrates ground truth (GT) annotations. The values below each plot indicate the spot-to-spot matching ratio.

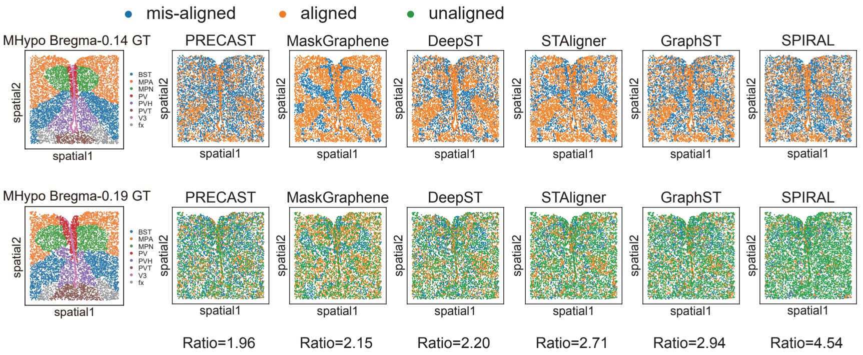

Supplementary Figure 2: **Visualization plots for alignment-misalignment-unalignment and spot-to-spot mapping ratio on MHypo Bregma -0.14 - -0.19 pair.** Visualization plots displaying aligned spots, misaligned spots, and unaligned spots during the alignment process. The anchor spot from the first (top) slice is aligned to the corresponding spots on the second (bottom) slice for MHypo Bregma -0.14 - -0.19 pair. The first slice pair illustrating ground truth (GT) annotations. The values below each plot indicate the spot-to-spot matching ratio.

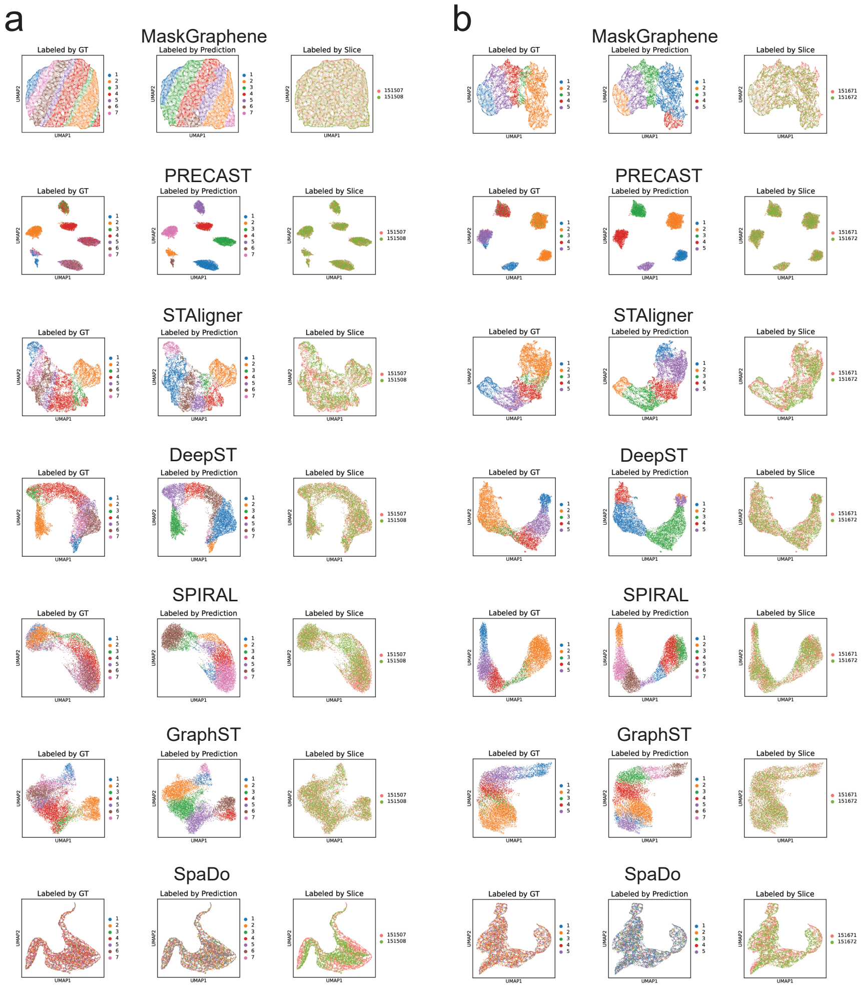

Supplementary Figure 3: **UMAP plots of low dimensional joint embedding on the DLPFC dataset.** **(a)** UMAP visualizations of joint embeddings generated by different methods for the DLPFC 151507-151508 pair-wise integration. Spots are colored by ground truth (GT) labels, predicted domains, and slice identity. Each row corresponds to a method: MaskGraphene, PRECAST, STAligner, DeepST, SPIRAL, GraphST, and SpaDo. **(b)** UMAP visualizations of joint embeddings generated by different methods for the DLPFC 151671-151672 pair-wise integration. Spots are colored by ground truth (GT) labels, predicted domains, and slice identity. Each row corresponds to a method: MaskGraphene, PRECAST, STAligner, DeepST, SPIRAL, GraphST, and SpaDo.

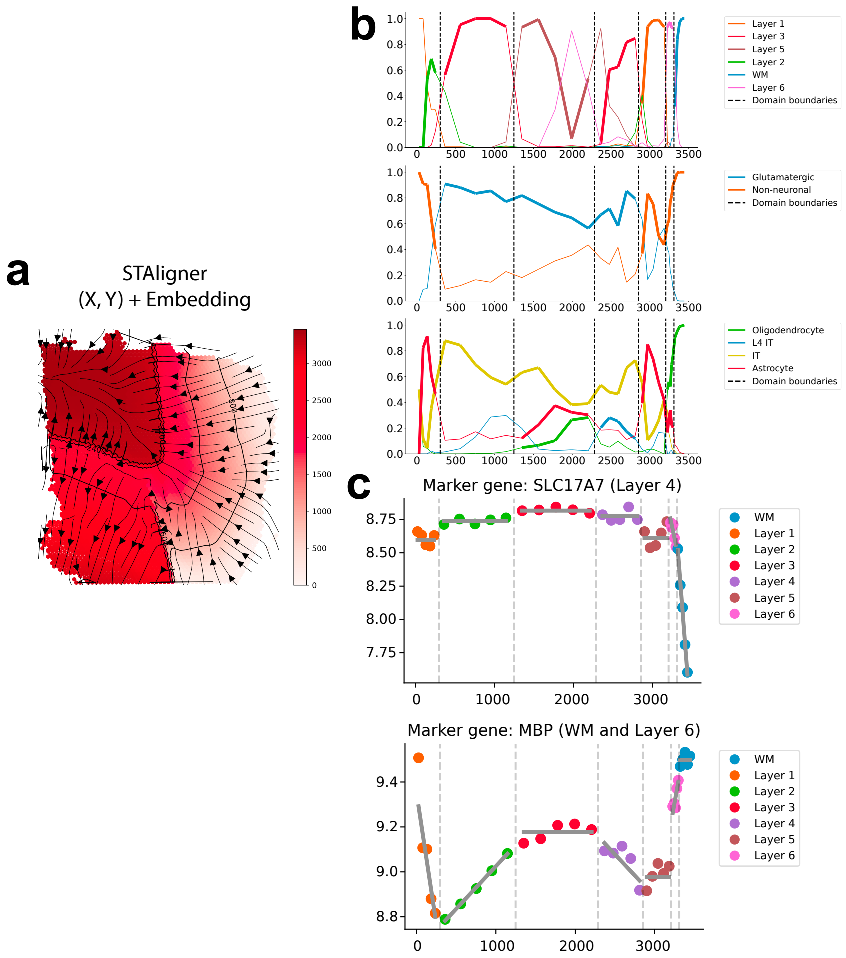

Supplementary Figure 4: **Topography analysis based on STAligner embeddings after DLPFC 151673-151674 pair-wise integration. (a)** Topographical maps generated using GASTON with original X,Y coordinates combined with joint embeddings of STAligner. **(b)** Plots showing the proportions of cell types as a function of the isodepth, using three different types of annotations: layer-specific cell types (top panel), neuronal types (middle panel), and cell types (bottom panel). **(c)** Plots showing the marker gene (SLC17A7 and MBP ) expression versus the isodepth.

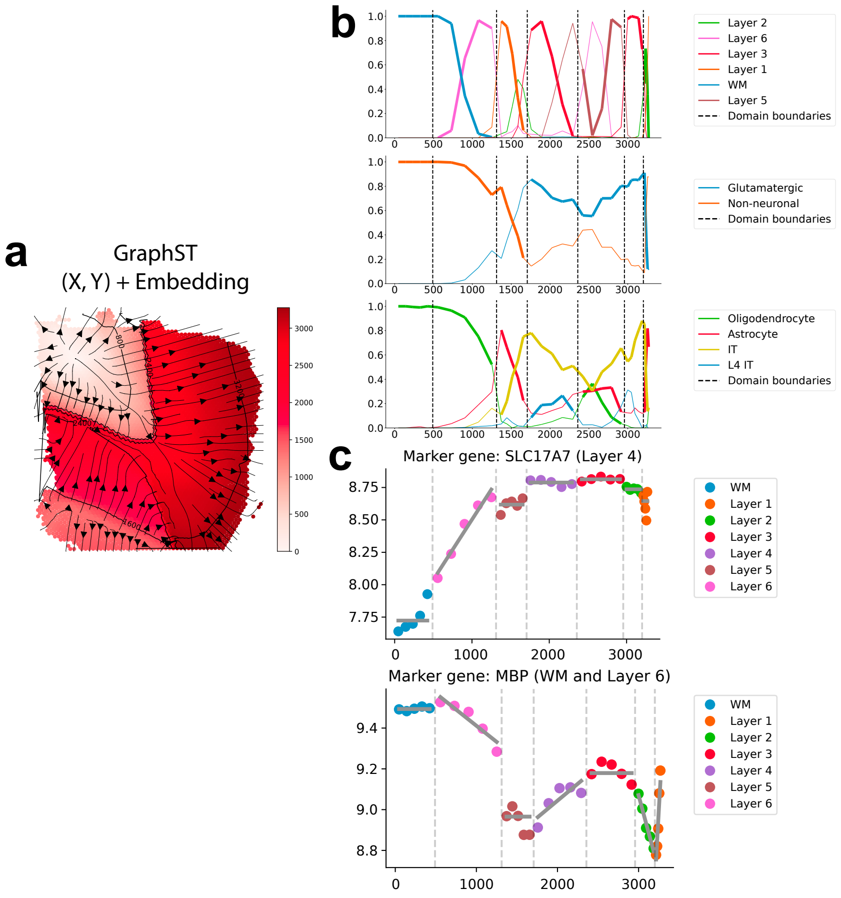

Supplementary Figure 5: **Topography analysis based on GraphST embeddings after DLPFC 151673-151674 pair-wise integration.** **(a)** Topographical maps generated using GASTON with original X,Y coordinates combined with joint embeddings of GraphST. **(b)** Plots showing the proportions of cell types as a function of the isodepth, using three different types of annotations: layer-specific cell types (top panel), neuronal types (middle panel), and cell types (bottom panel). **(c)** Plots showing the marker gene (SLC17A7 and MBP ) expression versus the isodepth.

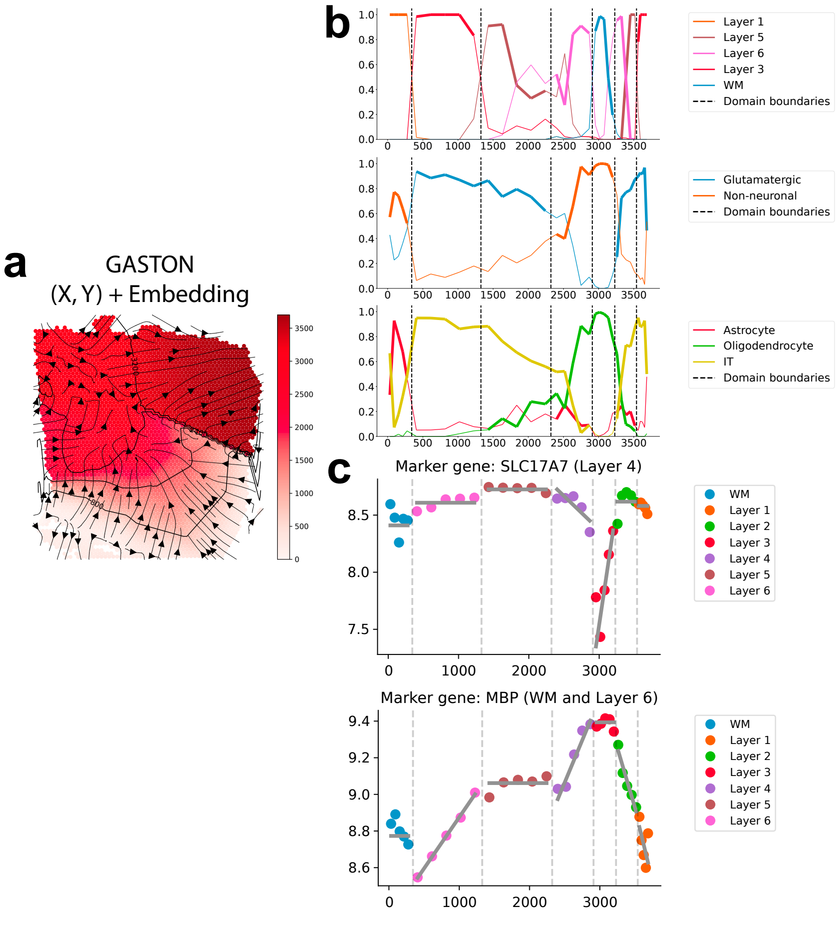

Supplementary Figure 6: **Topography analysis based on PCA-derived embeddings from individual DLPFC slice 151673.** **(a)** Topographical maps generated using GASTON with original X,Y coordinates combined with PCA-derived embeddings from DLPFC slice 151673 (GASTON's default setting). **(b)** Plots showing the proportions of cell types as a function of the isodepth, using three different types of annotations: layer-specific cell types (top panel), neuronal types (middle panel), and cell types (bottom panel). **(c)** Plots showing the marker gene (SLC17A7 and MBP ) expression versus the isodepth.

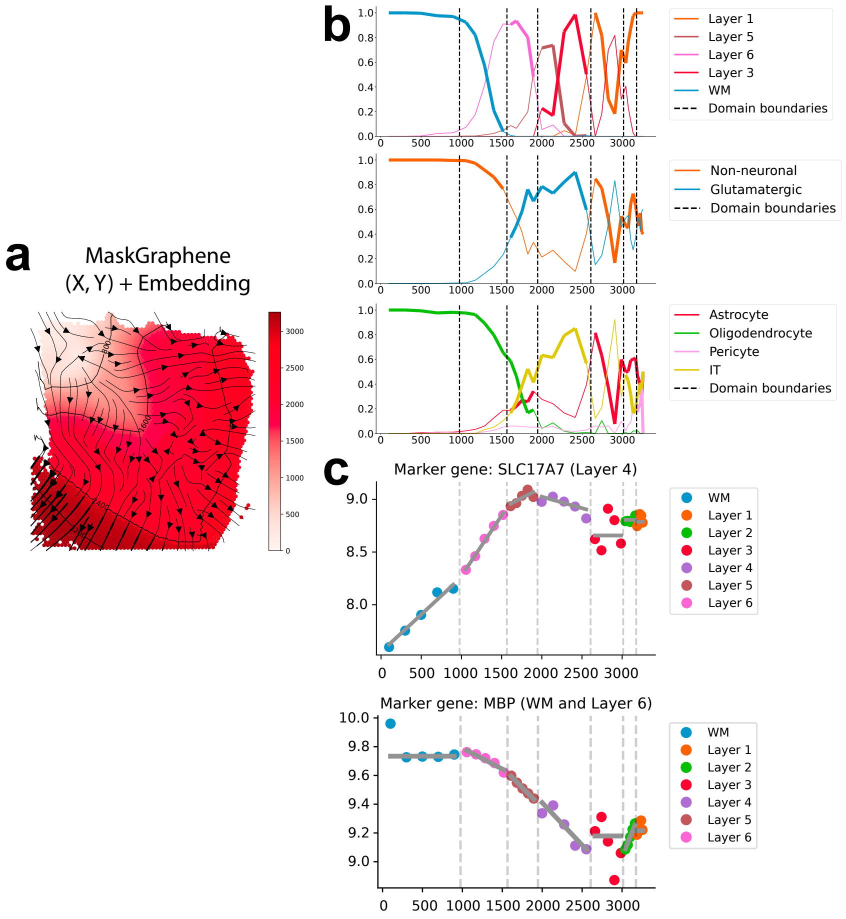

Supplementary Figure 7: **Topography analysis based on MaskGraphene embeddings after DLPFC four-slice integration (151673-151674-151675-151676).** **(a)** Topographical maps generated using GASTON with original X,Y coordinates combined with joint embeddings of MaskGraphene. **(b)** Plots showing the proportions of cell types as a function of the isodepth, using three different types of annotations: layer-specific cell types (top panel), neuronal types (middle panel), and cell types (bottom panel). **(c)** Plots showing the marker gene (SLC17A7 and MBP ) expression versus the isodepth.

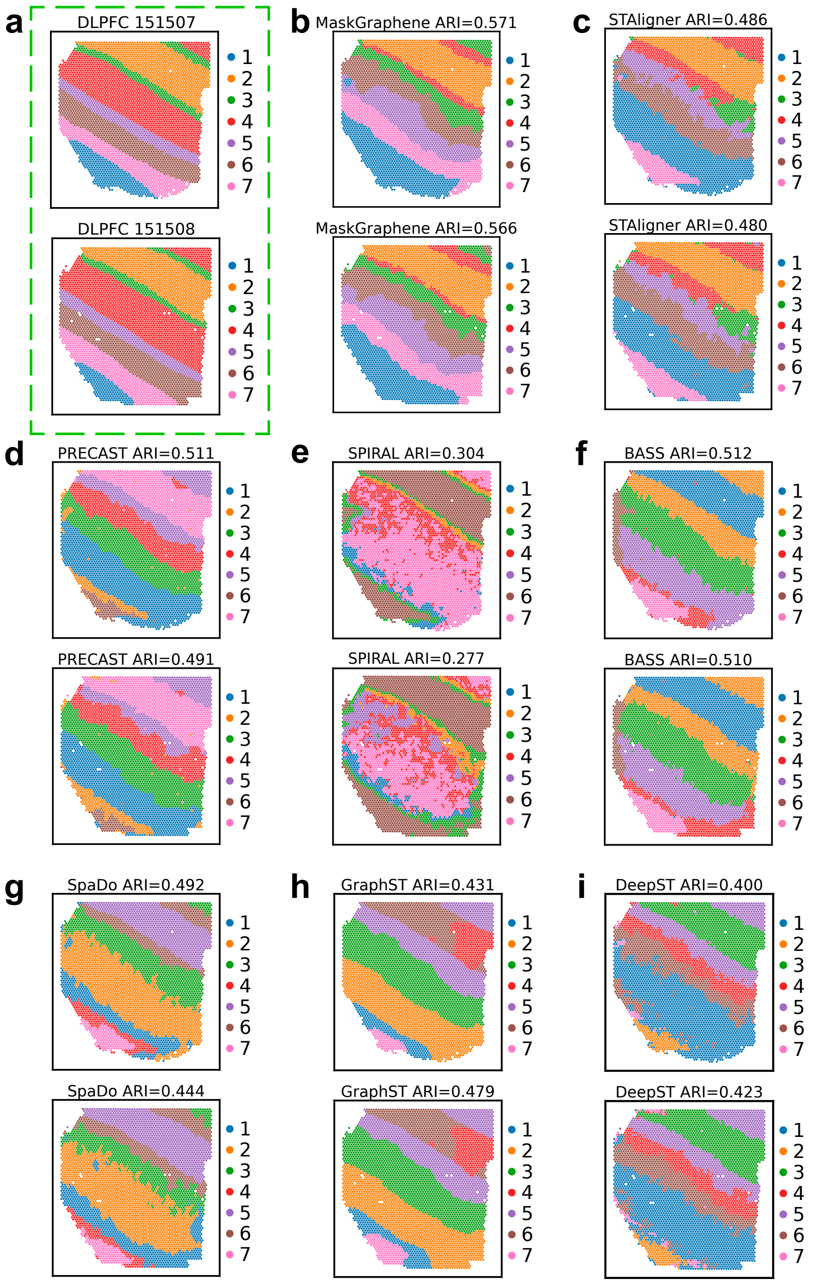

Supplementary Figure 8: **Spatial visualization of joint domain identification after DLPFC 151507-151508 pair-wise integration.** **(a)** Spatial domain identification visualization for two DLPFC slices (151507 and 151508) by ground truth. **(b-i)** Spatial domain identification visualization with ARI value for two DLPFC slices (151507 and 151508) after pair-wise integration using eight integration methods, including MaskGraphene (b), STAligner (c), PRECAST (d), SPIRAL (e), BASS (f), SpaDo (g), GraphST (h), and DeepST (i).

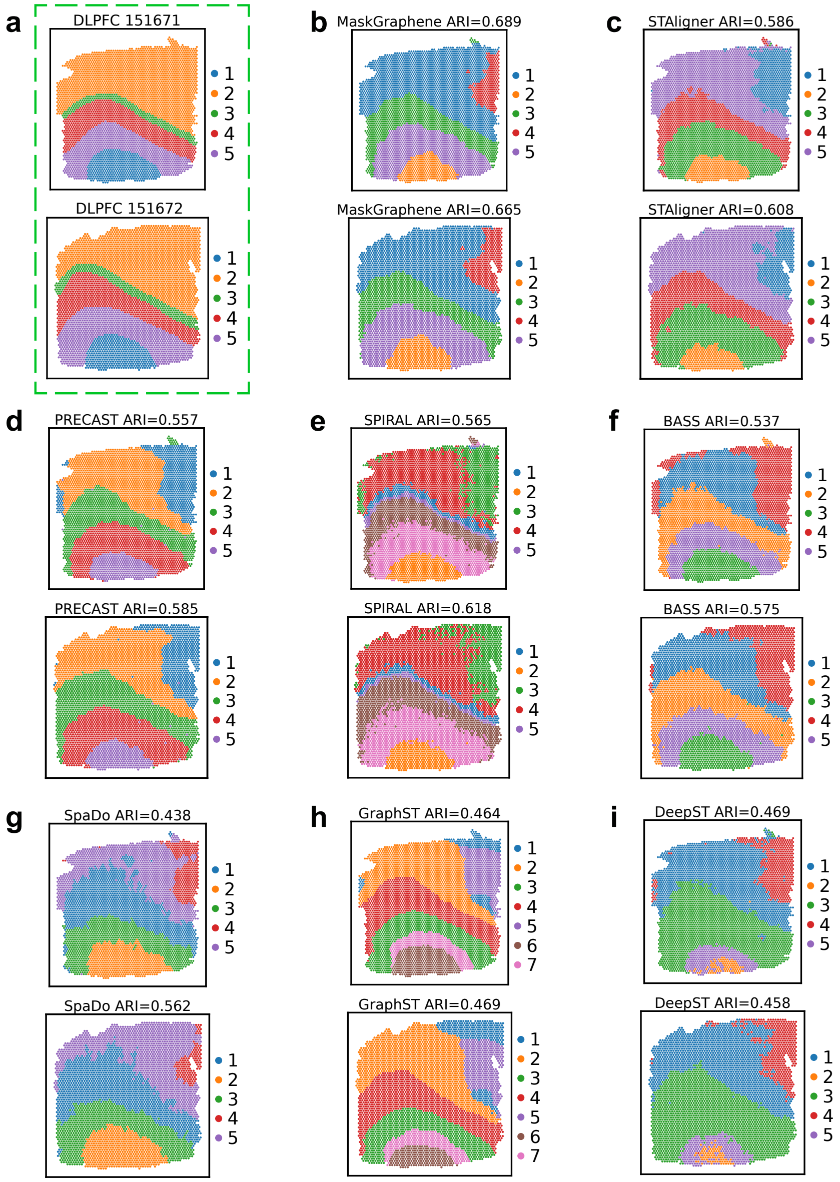

Supplementary Figure 9: **Spatial visualization of joint domain identification after DLPFC 151671-151672 pair-wise integration. (a)** Spatial domain identification visualization for two DLPFC slices (151671 and 151672) by ground truth. **(b-i)** Spatial domain identification visualization with ARI value for two DLPFC slices (151671 and 151672) after pair-wise integration using eight integration methods, including ground truth (a), MaskGraphene (b), STAligner (c), PRECAST (d), SPIRAL (e), BASS (f), SpaDo (g), GraphST (h), and DeepST (I).

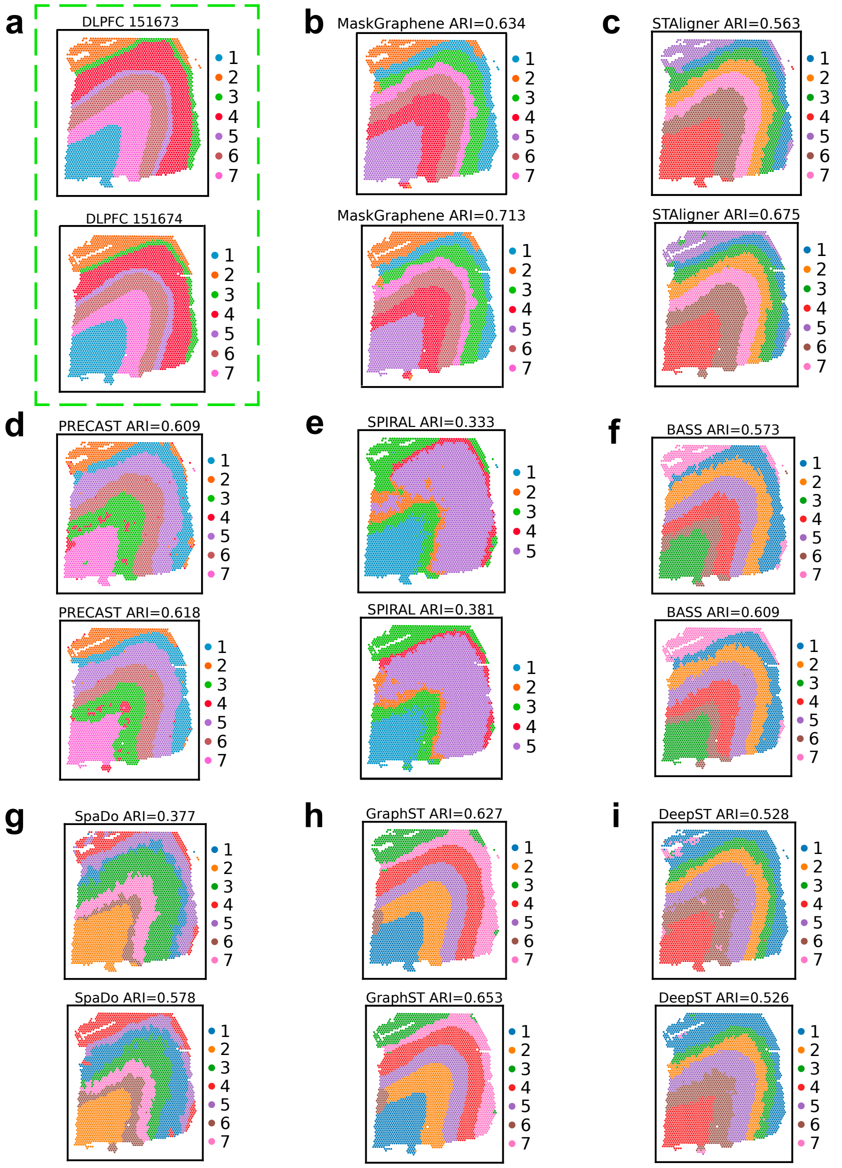

Supplementary Figure 10: **Spatial visualization of joint domain identification after DLPFC 151673-151674 pair-wise integration.** **(a)** Spatial domain identification visualization for two DLPFC slices (151673 and 151674) by ground truth. **(b-i)** Spatial domain identification visualization with ARI value for two DLPFC slices (151673 and 151674) after pair-wise integration using eight integration methods, including ground truth (a), MaskGraphene (b), STAligner (c), PRECAST (d), SPIRAL (e), BASS (f), SpaDo (g), GraphST (h), and DeepST (I).

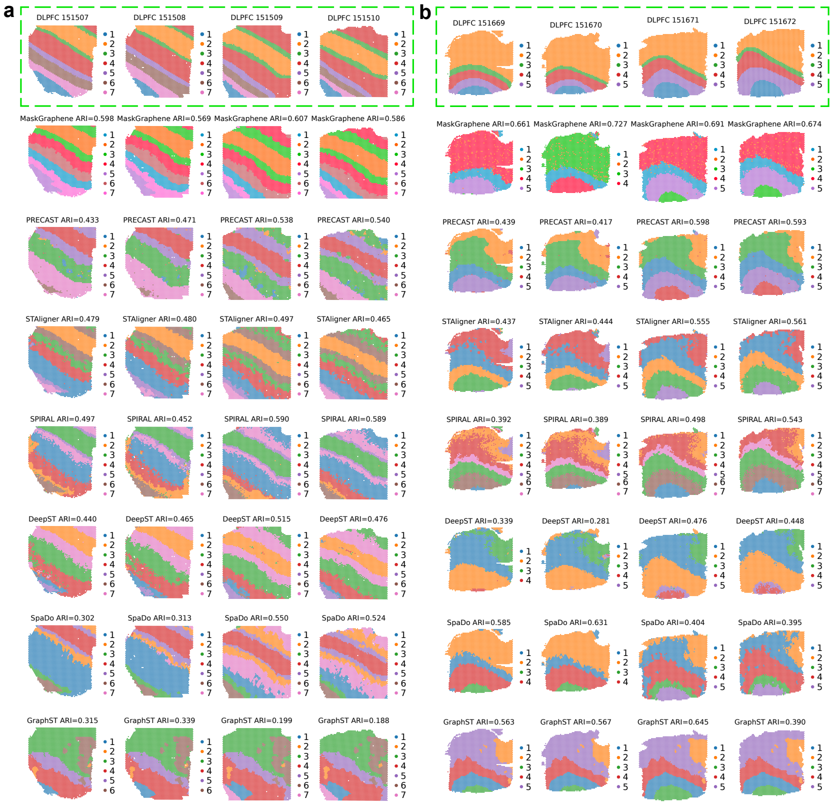

Supplementary Figure 11: **Spatial visualization of joint domain identification after four-slice integration (151507-151508-151509-151510).** **(a)** Spatial domain identification visualization with ARI value for four DLPFC slices (151507, 151508, 151509 and 151510) after four-slice integration using seven integration methods, including MaskGraphene, PRECAST, STAligner, SPIRAL, DeepST, SpaDo, and GraphST. Each row corresponds to a different method and ground truth. **(b)** Spatial domain identification visualization for four DLPFC slices (151669, 151670, 151671 and 151672) after integration using seven integration methods, including MaskGraphene, PRECAST, STAligner, SPIRAL, DeepST, SpaDo, and GraphST. Each row corresponds to a different method and ground truth.

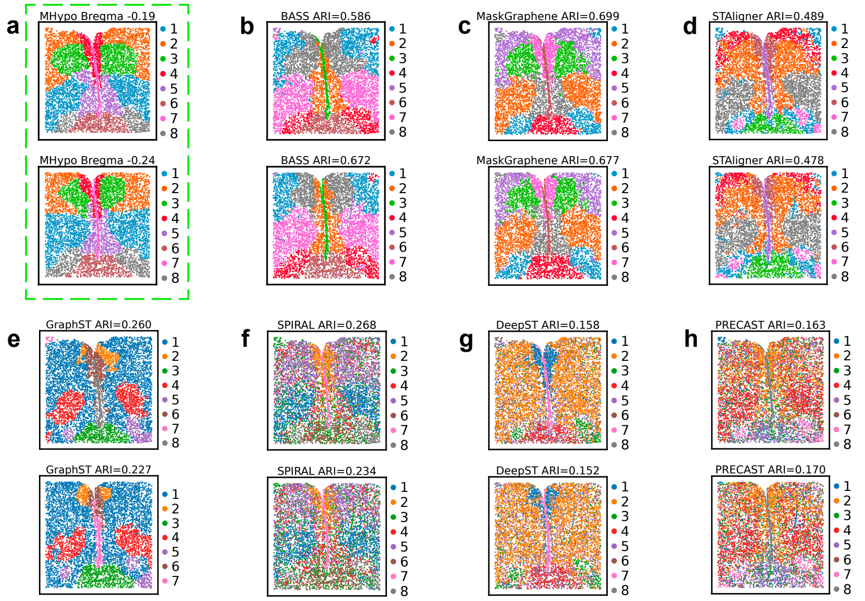

Supplementary Figure 12: **Spatial visualization of joint domain identification after MHypo Bregma -0.19 - -0.24 pair-wise integration. (a)** Spatial domain identification visualization for two MHypo slices (Bregma -0.19 and Bregma -0.24) by ground truth. **(b-h)** Spatial domain identification visualization with ARI value for two MHypo slices (Bregma -0.19 and Bregma -0.24) after pair-wise integration using seven integration methods, including BASS (b), MaskGraphene (c), STAligner (d), GraphST (e), SPIRAL (f), DeepST (g), and PRECAST (h).

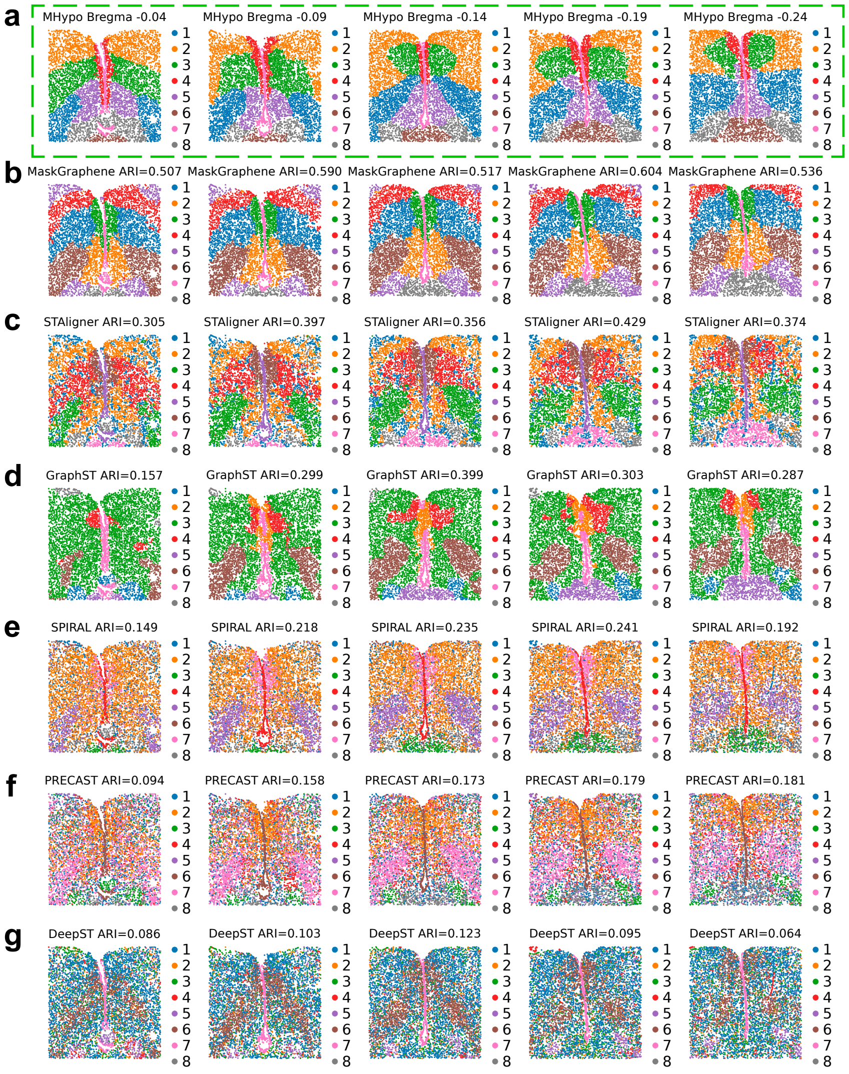

Supplementary Figure 13: **Spatial visualization of joint domain identification after MHypo five-slice integration. (a-g)** Spatial domain identification visualization for five MHypo slices (Bregma -0.04, Bregma -0.09, Bregma -0.14, Bregma -0.19 and Bregma -0.24) after five-slice integration using seven integration methods, including MaskGraphene, STAligner, GraphST, SPIRAL, PRECAST, and DeepST. Each row corresponds to a different method and ground truth.

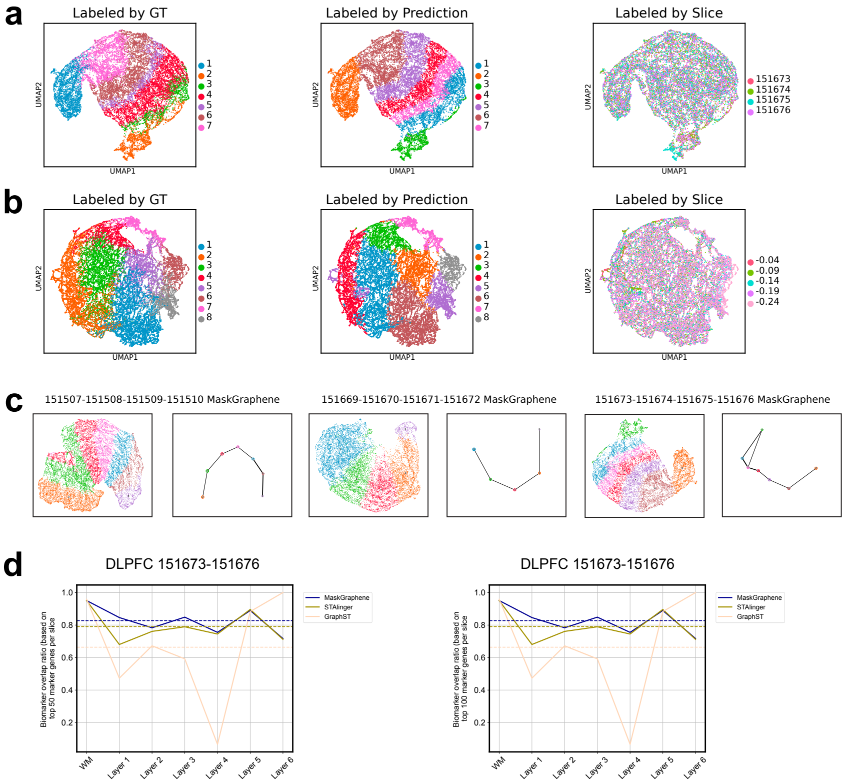

Supplementary Figure 14: **UMAP, PAGA, and biomarker analysis of MaskGraphene (coordinate transformation) after multi-slice integration.** **(a)** UMAP visualizations of joint embeddings generated by MaskGraphene (coordinate transformation) for the DLPFC four-slice integration (151673-151674-151675-151676). Spots are colored by ground truth (GT) labels, predicted domains, and slice identity. This subfigure corresponds to Figure 4b using MaskGraphene (coordinate replacement). **(b)** UMAP visualizations of joint embeddings generated by MaskGraphene (coordinate transformation) for the MHypo five-slice integration (Brega -0.04 - -0.09 - -0.14 - -0.19 - -0.24). Spots are colored by ground truth (GT) labels, predicted domains, and slice identity. This subfigure corresponds to Figure 5b using MaskGraphene (coordinate replacement). **(c)** Each two panels shows UMAP visualizations paired with PAGA graphs by MaskGraphene (coordinate transformation), illustrating spatial trajectory results for three distinct DLPFC four-slice integration: (151507-151508-151509-151510), (151669-151670-151671-151672), and (151673-151674-151675-151676). Spots are colored according to predicted domains. This subfigure corresponds to Figure 6 using MaskGraphene (coordinate replacement). **(d)** Biomarker overlap ratio curves across WM and cortical layers for MaskGraphene (coordinate transformation), comparing the top $N$ biomarkers identified for each integrated layer after four-slice integration with the $N$ ground truth marker genes. Ground truth markers are defined as the union of the top 50 (left panel) or top 100 (right panel) layer marker genes across four slices. This subfigure corresponds to Figure 7b using MaskGraphene (coordinate replacement).

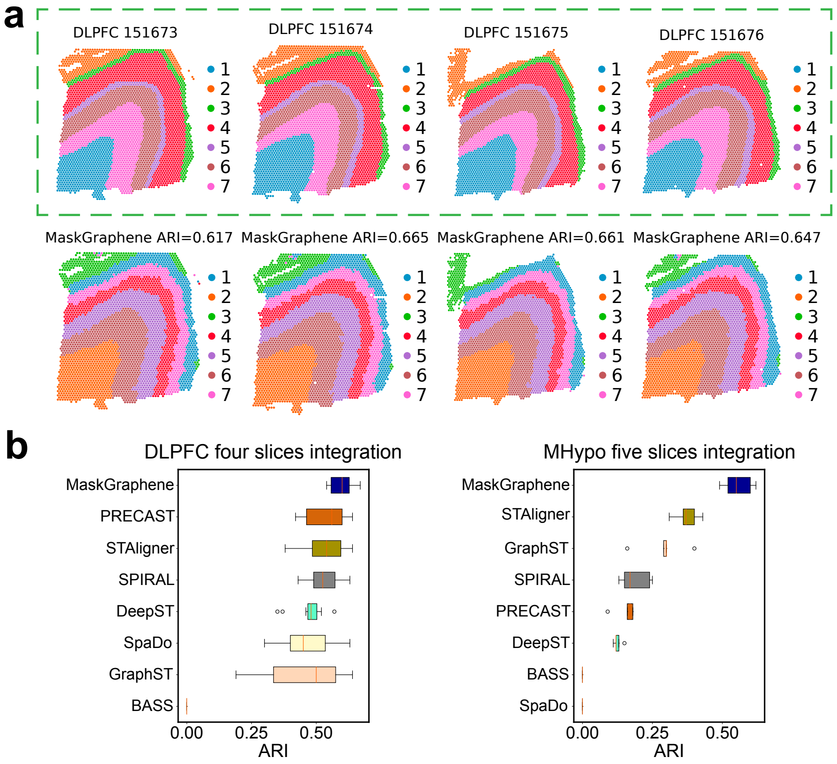

Supplementary Figure 15: **Spatial visualization of joint domain identification and ARI boxplots by MaskGraphene (coordinate transformation) after multi-slice integration. (a)** Visualization of spatial domain identification for four DLPFC slices (151673, 151674, 151675, and 151676), showing ground truth (top panels enclosed in a green dashed box) and results after integration using MaskGraphene (coordinate transformation) (bottom panels) **(b)** Box plots showing ARI scores for all DLPFC four-slice integration and all MHypo five-slice integration by MaskGraphene (coordinate transformation) and all other integration methods. This Figure corresponds to Figure 9 using MaskGraphene (coordinate replacement).

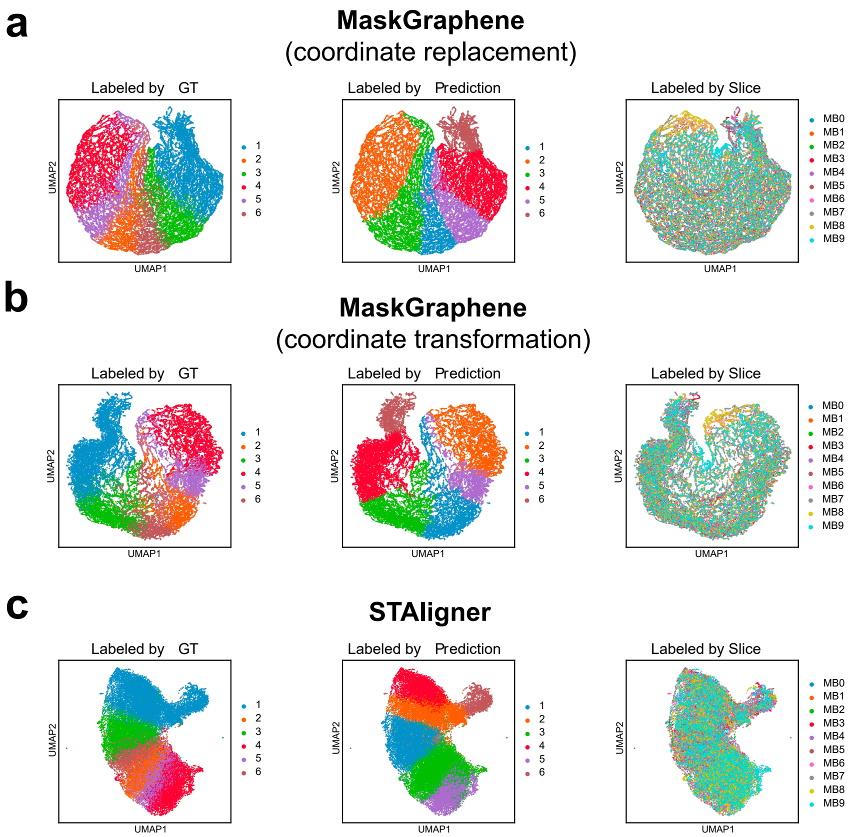

Supplementary Figure 16: **UMAP plots of low dimensional joint embedding on the MB dataset.** **(a)** UMAP visualizations of joint embeddings generated by MaskGraphene (coordinate replacement). Spots are colored by ground truth (GT) labels, predicted domains, and slice identity. **(b)** UMAP visualizations of joint embeddings generated by MaskGraphene (coordinate transformation). Spots are colored by ground truth (GT) labels, predicted domains, and slice identity.**(b)** UMAP visualizations of joint embeddings generated by STAligner. Spots are colored by ground truth (GT) labels, predicted domains, and slice identity.

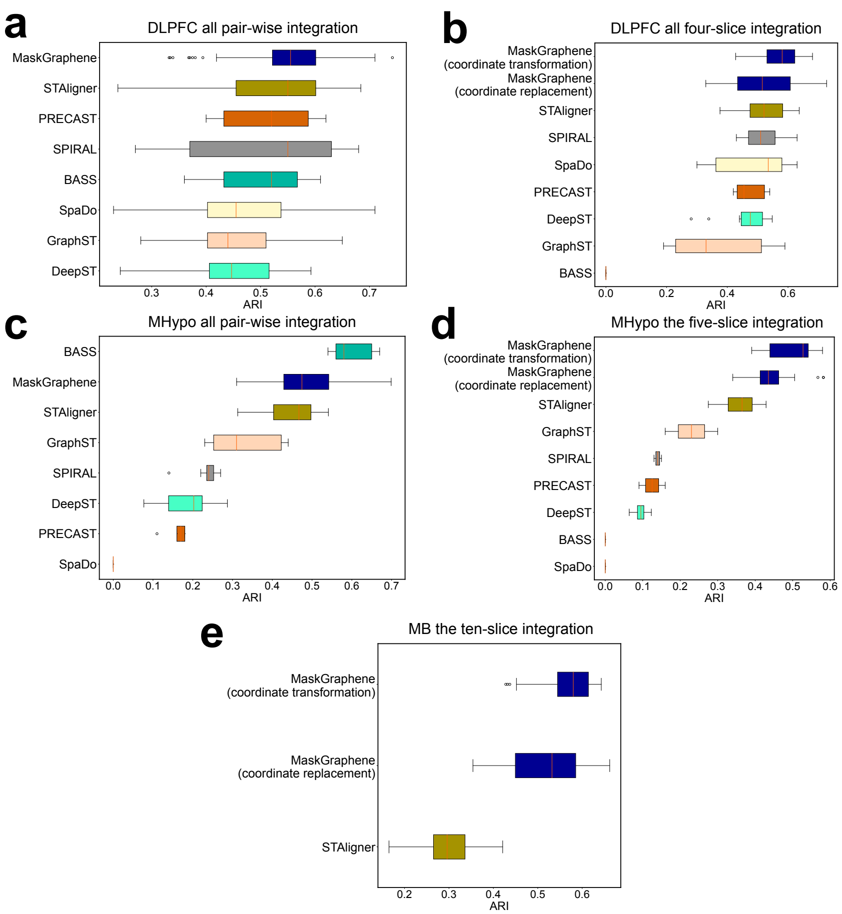

Supplementary Figure 17: **ARI box plots of clustering results after integration with varying random seeds across different datasets and methods.** **(a)** ARI box plots of clustering results after all DLPFC pair-wise integration with varying random seeds. **(b)** ARI box plots of clustering results after all DLPFC four-slice integration with varying random seeds. **(c)** ARI box plots of clustering results after all MHypo pair-wise integration with varying random seeds. **(d)** ARI box plots of clustering results after the MHypo five-slice integration with varying random seeds. **(e)** ARI box plots of clustering results after the MB ten-slice integration with varying random seeds.
